# Argonaute Binding within 3’-Untranslated Regions Does Not Predict Gene Repression

**DOI:** 10.1101/2020.02.25.956623

**Authors:** Yongjun Chu, Audrius Kilikevicius, Jing Liu, Krystal C. Johnson, Shinnichi Yakota, David R. Corey

## Abstract

Despite two decades of study, the full scope of RNAi in mammalian cells has remained obscure. Here we combine: 1) Knockout of argonaute (AGO) variants; 2) RNA sequencing analysis of gene expression changes; and 3) Crosslinking Immunoprecipitation Sequencing (CLIP-seq) using anti-AGO2 antibody to identify potential microRNA (miRNA) binding sites. We find that knocking out AGO1, AGO2, and AGO3 are necessary to achieve full impact on gene expression. CLIP-seq reveals several hundred significant AGO2 associations within the 3’-untranslated regions of cytoplasmic transcripts. The standard mechanism of miRNA action would suggest that these associations repress gene expression. Contrary to this expectation, clusters are poorly correlated with gene repression in wild-type versus knockout cells. Many clusters are associated with increased gene expression in wild-type versus knock out cells, including the strongest cluster within the *MYC* 3’-UTR. Our results suggest that assumptions about miRNA action should be re-examined.

## INTRODUCTION

RNA interference is a powerful mechanism for controlling gene expression in mammalian cells (Hammond, 2015; Bartel, 2018) and is now achieving success in the clinic (Shen and Corey, 2018; Setten et al. 2019). Chromosomally-encoded hairpin RNAs are expressed and are processed into microRNAs (miRNAs) in the cytoplasm (Michlewski and Caceres, 2019). These miRNAs are loaded into argonaute protein (AGO) to from a ribonucleoprotein in which the miRNA programs the complex to bind complementary RNA sequences while AGO increases the efficiency of binding and allows association with GW182 and other proteins to enhance function (Meister, 2013; Yao et al., 2013). miRNAs are thought to require only partial complementarity to target sequences, with a match at bases 2-8 being most important for recognition (Lewis et al. 2003; Jackson et al., 2006).

There are four AGO proteins in human cells, AGO1-4 (Meister 2013). AGO2 is the best studied and is known as the catalytic engine for RNAi because it possesses an enzyme active site capable of cleaving target RNA transcripts (Liu et al. 2004, Meister et al. 2004). This ability to cleave mRNA efficiently is critical for application of fully complementary synthetic RNAs to gene silencing and contributes to the robustness of RNAi as a method for controlling gene expression in the laboratory and the clinic. The roles of AGO1 and AGO3 are less known. One report has suggested that each AGO is functionally equivalent with bulged miRNA duplexes during the action of miRNAs, while AGO1 and AGO2 may be more active in conjunction with duplex RNAs (Su et al., 2009).

Since 2000, a literature search of the term miRNA reveals over 90,000 citations with over 10,000 new citations appearing every year. 42,000 papers appear on a PubMed search of “cancer” and “miRNA”. These numbers suggest that the science of miRNA action is well settled. The assumption that miRNAs should have widespread roles in regulating cell biology has been based on several factors: 1) Efficient RNAi gene regulation in *C. elegans* and other model organisms (Fire et al., 1998; Zamore et al., 2000; Elbashir et al., 2001a); 2) the simplicity of base-pairing rules that encourage hypotheses connecting a miRNA seed sequence to a potential target gene of interest; and 3) the known robustness of fully complementary synthetic RNAs as gene silencing agents in mammalian cells (Elbashir et al, 2001b).

Closer examination, however, reveals that many papers lack the minimum controls and experimentation necessary to make convincing conclusions linking complementary recognition by a miRNA to a functional effect on gene expression (Gagnon and Corey, 2019). One possible reason for the proliferation of these studies claiming diverse and often conflicting regulatory roles for miRNAs is an inadequate understanding of the quantitative and biochemical foundations for RNAi and miRNAs inside human cells.

We have used HCT116 cells to investigate RNAi and have compared gene expression in AGO1, AGO2, AGO1/2, and AGO1/2/3 knock out cells to wild-type cells. Knocking out AGO1, AGO2, and AGO3 was necessary to achieve full effects. We used enhanced crosslinking immunoprecipitation to identify locations for AGO binding with 3’-untranslated regions (3’-UTRs), the accepted target for repression by miRNAs. Contrary to expectations, we observe no correlation between AGO binding within the 3’-UTR gene repression – most genes with clusters showed no change in expression or decreased expression upon knockout of one or more AGO variants. The lack of correlation between the association of AGO2 within 3’-UTRs and gene repression suggests reexamination of previous assumptions about miRNA action.

## MATERIALS AND METHODS

### Cell lines

The HCT116 cell line (Horizon Discovery) originated from the American Type Culture Collection (ATCC) and then licensed and supplied to the European Collection of Authenticated Cell Cultures (ECACC). ATCC authenticated this HCT116 cell line using Short Tandem Repeat (STR) analysis as described in 2012 ANSI Standard (ASN-0002) Authentication of Human Cell Lines: Standardization of STR profiling by the ATCC Standards Development Organization (SDO) (Capes-Davis et al., 2012). The ATCC STR analysis compared seventeen short tandem repeat loci plus the gender determining locus, Amelogenin, to verify the HCT116 cell line (ATCC CCL 247). ECACC performed an additional STR analysis of seventeen loci on the cells received from ATCC, and the verified HCT116 cells (ECACC 91091005) were supplied to Horizon Discovery.

For eCLIP, we used HCT116:AGO2 knockout cells obtained from Joshua T. Mendell (UT Southwestern). The AGO1, AGO2, AGO1/2 and AGO1/2/3 knockout cell lines used for RNAseq were prepared using GenCRISPR™ gene editing technology and services (GenScript) and verified HCT116 cells (Horizon Discovery). The AGO2 knockout cell line was independently generated to ensure that all cell lines used for RNAseq were derived from similar genetic backgrounds. gRNA sequence, gRNA target locations, and DNA sequencing results are shown in Figure 2. All HCT116 and HCT116-derived cells were cultured in McCoy’s 5A Medium (Sigma-Aldrich) supplemented with final 10% FBS at 37 °C in 5% CO_2_. For the cell growth assay, the cells were seeded at a density of 50,000 cells/mL, disassociated with 1X trypsin, and counted using using trypan blue staining (TC20 Automated Cell Counter, Bio-Rad).

### Preparation of cytoplasmic extract

Cytoplasm isolation was similar as previously described (Gagnon et al. 2014A;Gagnon et al. 2014B), with modifications for HCT116 and HCT116-derived cells. Cells at ∼95% confluence were lysed in hypotonic lysis buffer (HLB) (10 mM Tris-HCl, pH-7.4, 10 mM NaCl, 3 mM MgCl_2_, 2.5% NP-40, 0.5 mM DTT, 1x protease inhibitor (Roche, cOmplete) and 50 U/mL ribonuclease inhibitors (RNasin^®^ Plus, Promega) and supernatant collected as cytoplasmic fraction. Western blots to determine the purity of fractions and RNAi factors distribution were performed as before (Gagnon et al. 2014) using anti-AGOs (defined above) **SPECIFY**, anti-GW182, 1:5000 (A302-329A, Bethyl Laboratories), anti-β-tubulin, 1:5000 (T5201, Sigma-Aldrich), anti-Calnexin, 1:1000 (2433S, Cell Signaling), anti-LaminA/C, 1:1500 (ab8984, Abcam) and anti-Histone 3, 1:20000 (2650S, Cell Signaling).

### RT-qPCR

Total RNA was extracted from HCT116 wild-type or knockout cells and treated with DNase I (Worthington Biochemical) at 25 °C for 20 min, 75 °C for 10 min. Reverse transcription was performed using high-capacity reverse transcription kit (Applied Biosystems) per the manufacturer’s protocol. 2.0 µg of total RNA was used per 20 µl of reaction mixture. PCR was performed on a 7500 real-time PCR system (Applied Biosystems) using iTaq SYBR Green Supermix (BioRad). PCR reactions were done in triplicates at 55 °C 2 min, 95 °C 3 min and 95 °C 30 s, 60 °C 30 s for 40 cycles in an optical 96-well plate. For each gene, two different sets of primer are used to check mRNA level (**Supplementary Table S1**). Data were normalized relative to measured hypoxanthine phosphoribosyltransferase 1 (HPRT1) and Small Nuclear Ribonucleoprotein U5 Subunit 200 (snRNP200) genes level.

### RNAseq for gene expression analysis

WT HCT116, AGO1, AGO2, AGO1/2, and AGO1/2/3 knock out cells were used for RNAseq. Three biological replicated samples were sequenced. Approximately 3.0×10^6^ cells were seeded in a 15-cm large dish. Cells were harvested 48 hours later and RNA was extracted using the RNeasy Mini Kit (Qiagen) with an on-column DNase digestion. Sequencing libraries were generated using the TruSeq Stranded Total RNA with Ribo-Zero Human/ Mouse/Rat Low-throughput (LT) kit (Illumina) and run on a NextSeq 500 for paired-end sequencing using the NextSeq 500/550 High Output v2 Kit, 150 cycles (Illumina).

Quality assessment of the RNA-seq data was done using NGS-QC-Toolkit43 with default settings. Quality-filtered reads generated by the tool were then aligned to the human reference genome hg38 and transcriptome gencode v75 using the STAR (v 2.5.2b) using default settings. Read counts obtained from STAR were used as input for Salmon (v 1.0.0) and Deseq2 for gene differential expression analysis. Genes with adjusted *p* ≤ 0.05 were regarded as differentially expressed for comparisons of each sample group.

### Enhanced UV crosslinking and immunoprecipitation (eCLIP)

#### Sequencing library prep

Control and AGO2 −/− HCT116 cells (obtained from Dr. Joshua Mendel, UT Southwestern) were seeded in 15 cm dishes with 12 dishes per cell line at 3.0×10^6^ cells per dish. Cells were cultured for 48 hours and subsequently UV crosslinked at 300 mJ/cm^2^. Cytoplasmic fraction was collected as described above. eCLIP was performed using the frozen samples as previously described (Van Nostrand et al., 2016), using anti-AGO2 antibody for IPs (3148, gift from Jay A. Nelson lab). For each cell line, duplicate input and IP samples were prepared and sequenced. The RiL19 RNA adapter (Supplementary Table 1) was used as the 3’ RNA linker for all samples. PAGE purified DNA oligonucleotides were obtained from IDT for the PCR library amplification step (Supplementary Table 1). PCR amplification was performed using between 11-16 cycles for all samples. Paired-end sequencing was performed on a NextSeq 500 using the NextSeq 500/550 High Output v2 Kit, 100 cycle (Illumina).

#### Mapping deep sequencing reads

Adapters were trimmed from original reads using Cutadapt (v1.9.1) with default settings. Next, the randomer sequence from the rand103Tr3 linker (Supplementary Table 1) was trimmed and recorded. STAR (v2.5.2b) was used to align mate 2 to hg38. Only the uniquely mapped reads were retained. PCR duplicates were then removed using the randomer information using an in-house script. All reads remaining after PCR duplicate removal were regarded as usable reads and used for cluster calling.

#### eCLIP cluster calling and annotation

eCLIP clusters were identified following a method described by original eCLIP developers (VanNostrand et al., 2016) with the following modifications. Initial AGO2 binding clusters identified by CLIPper in wild-type HCT116 were filtered to keep only clusters that are statistically significant (p < 0.001). For each region, normalization to total usable reads was performed and a fold change between IP and combined samples (input and IP in knockout cell line samples) was calculated. Significant CLIP clusters in each dataset were defined by i) the presence of significantly greater coverage in the region than expected by chance based on the Poisson distribution (see above), and ii) log2 fold change of normalized reads in the cluster was ≥ 2 comparing IP to combined (input + IP in knockout cells).

The final CLIP clusters for AGO2 were identified by first identifying significant clusters present in both experimental replicates. A cluster was considered to be present in both replicates if it occurred on the same strand and the replicate clusters overlapped by at least 1/3 of their total length. Significant clusters from both replicates were then merged to define the final cluster length. Clusters were annotated based on their genomic locations (Gencode v27). If a cluster was assigned to multiple annotations, the annotation was selected using the following priority: CDS exon > 3’ UTR > 5’ UTR > Protein-coding gene intron > Noncoding RNA exon > Noncoding RNA intron > Intergenic.

## RESULTS

### Knockout of AGO1, AGO2, and AGO3 in HCT116 cells

We obtained CRISPR gene knockouts to investigate the role of AGO in controlling gene expression. HCT116 cells were chosen as a parental line because they are diploid, simplifying the process of obtaining CRISPR-mediated double or triple gene knockouts. HCT116 is a typical cell line in terms of miRNA expression (Ghandi et al., 2009) (**Supplementary Figure 1**). The AGO1, AGO2, AGO1/2, and AGO1/2/3 cells were verified by DNA sequencing (**Figure 1A-C**). Western analysis confirmed that the knock out cell lines did not express the targeted proteins (**Figure 1D**). AGO4 KO cells were not obtained because we had previously used quantitative mass spectrometry to determine that AGO4 was expressed at barely detectable levels in wild-type HCT116 cells (Liu et al., 2019).

**Figure 1.**
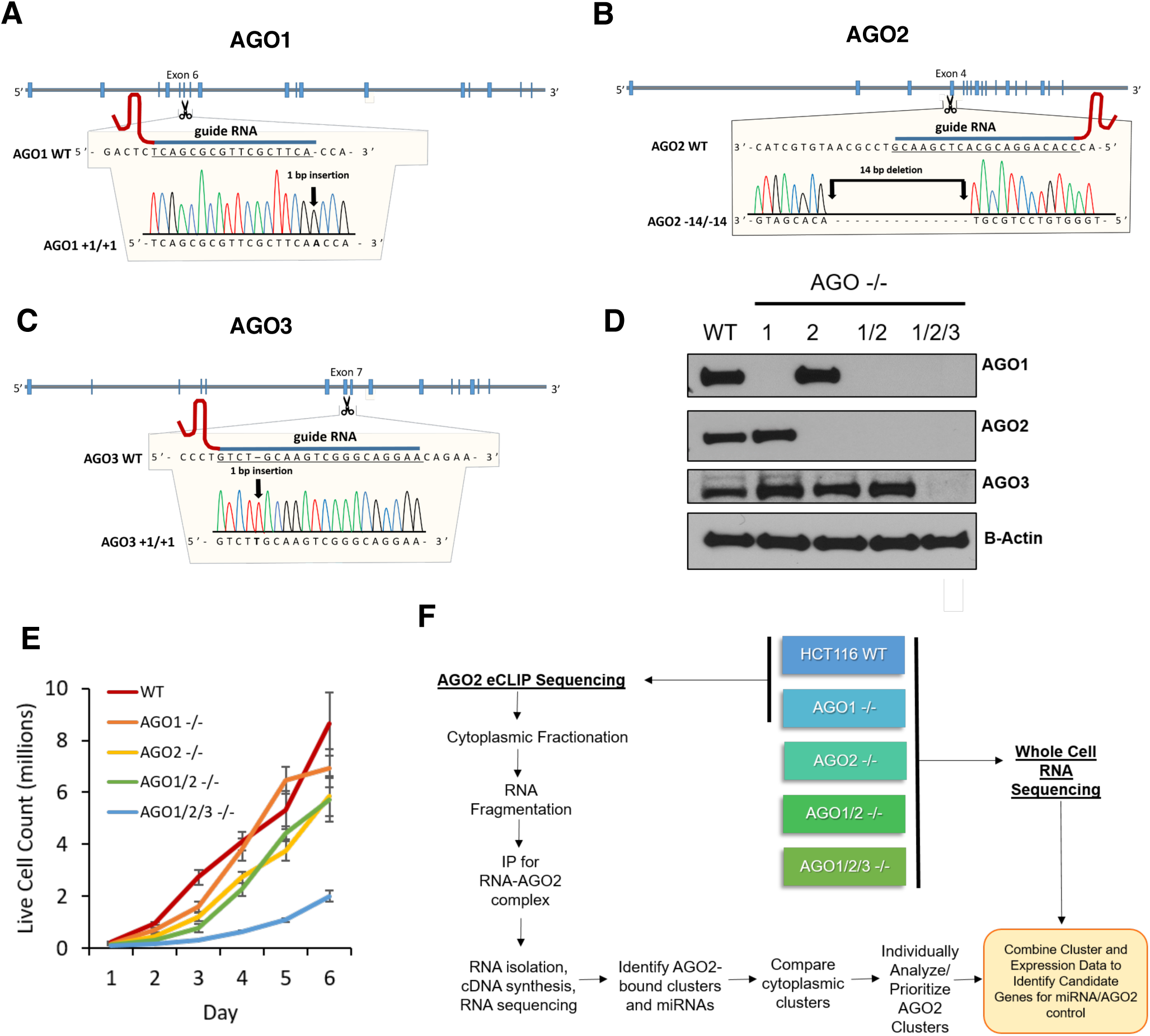
Characterization of CRISPR-derived HCT116AGO1/2/3 knockout cell lines. (**A-C**) Location of guide RNA and characterization of mutations for knocking out AGO1, AGO2, and AGO3. (**D**) Western blot validating the AGO1, AGO2, AGO1/2, and AGO1/2/3 knockout cell lines (n=3). **(E**) HCT116 WT, AGO1, AGO2, AGO1/2, and AGO1/2/3 knockout cell lines were seeded at the same density, harvested, and analyzed for live cell count (n=3). (**F**) Scheme showing knockout of AGO variants, RNAseq analysis of gene expression changes relative to wild-type cells; and CLIP-sequencing with anti-AGO2 antibody to identify potential miRNA-AGO2 binding sites. See also Figure S1.

We evaluated the growth rates of the knockout cell lines relative to wild-type (**Figure 1E**). The growth rates of AGO1 knockout and wild-type were similar. AGO2 and AGO1/2 knockout cells grew significantly more slowly than wild-type cells. The AGO1/2/3 triple knock out cells grew at much slower rates, consistent with the suggestion that knocking out all three primary AGO variants was necessary to achieve a greater physiologic effect than could be achieved in the single or double AGO knockout cells.

Enhanced Crosslinking Immunoprecipitation (eCLIP) RNAseq is a version of CLIP designed to increase signal to noise and improve confidence in the detection of potential sites of protein:RNA association (Van Nostrand et al., 2016) (**Figure 1F**). To determine sites of AGO binding we used an anti-AGO2 antibody. We focused on AGO2 because it is the best studied AGO variant and because the anti-AGO2 antibody was well-characterized as efficient for pull-down experiments (Kalantari et al. 2016). We used RNAseq to measure whole cell RNA levels in wild-type and knockout HCT116 cells for all AGO knockout cell lines. The correlation of AGO-binding data and gene expression data allowed us to classify the effect on AGO knockouts on the expression of genes with significant clusters.

### Distribution of AGO2 binding clusters

For eCLIP, we isolated RNA from the cytoplasmic fraction of HCT116 cells (**Supplementary Figure S2**). Parallel experiments were performed on HCT116 cells with AGO2 gene expression knocked out to subtract background signal. The significance of clusters was calculated using the Fisher exact test which considers the combined signals from Input and AGO2-eCLIP with AGO2 KO cells. A p-value less than 0.05 and an enrichment fold over 4 for wild-type versus AGO2 knockout cells were applied as threshold for a peak to be called statistically significant. All clusters were individually curated and visually examined to gain insights into biological significance and potential mechanisms of action.

Consistent with previous results (Chi et al., 2009; Hafner et al., 2010; Erhard et al., 2013; Moore et al., 2014) the largest category of AGO2 binding clusters detected by eCLIP were within the 3’-UTR of genes (**Figure 2A**). In many cases, multiple clusters were within the same gene and 1077 genes possessed significant clusters within their 3’-UTRs. The strongest (most significant) clusters were within miRNAs including 90% of the top two hundred clusters (**Figure 2BC**). Strong association with miRNAs may be because AGO2 binds directly to miRNA and because miRNAs tend to be much more highly expressed than mRNAs (Bosson et al., 2014). Of the top 200 clusters that were not miRNAs, 50% were within the 3’-UTR (**Figure 2D**). Examination of the top 250 clusters within the 3’-UTR revealed that clusters are distributed evenly throughout the region (**Figure 2E**).

**Figure 2.**
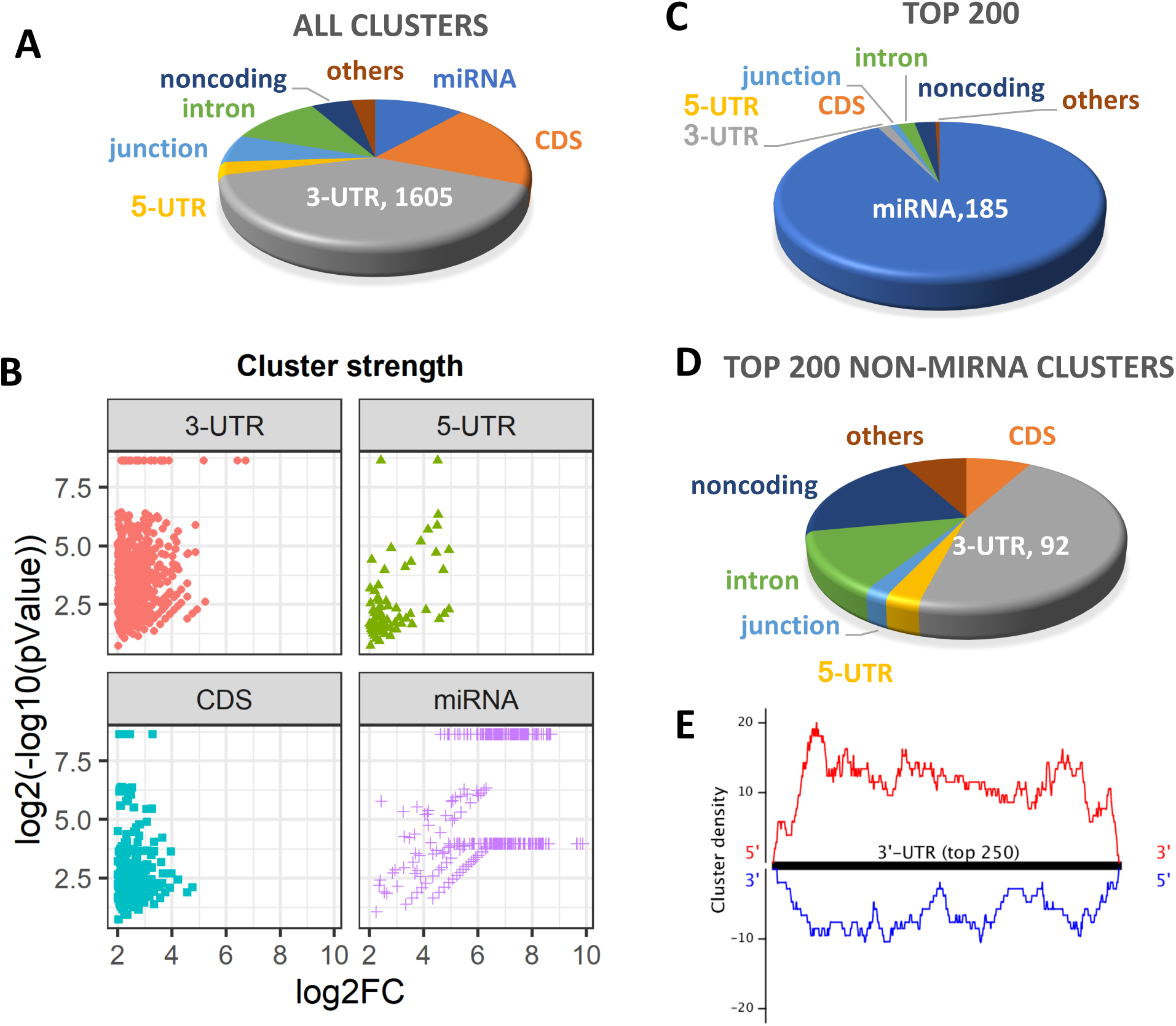
Localization of AGO2 binding clusters disclosed by eCLIP. (**A**) Relative distribution of all clusters. (**B**) Strength of clusters with different transcript regions. (**C,D**). Relative distribution of of top clusters, ranked by significance for the top 200 clusters and the top 200 clusters that that are not directly associated with miRNAs. (**E**) Localization of the strongest 250 clusters within 3’-UTRs. See also Figure S2.

### miRNA expression in HCT116 cells

We used our eCLIP data to estimate the relative abundance of miRNAs in cytoplasm of HCT116 cells (**Figure 3A**). Seventeen miRNAs had over 1000 reads, accounting for >70% of all reads that originate from miRNAs. Half of all reads were associated with just six miRNA families (**Figure 3B**). Visual examination of read clusters for top ranked miRNAs confirms unambiguous association of AGO2 and clear discrimination relative to AGO knock out cells and input controls (**Supplemental Figure S3**). Quantitation of miRNA abundance using qPCR revealed that highly expressed miRNAs were present at 1000-3000 copies per cell (**Figure 3C**).

**Figure 3.**
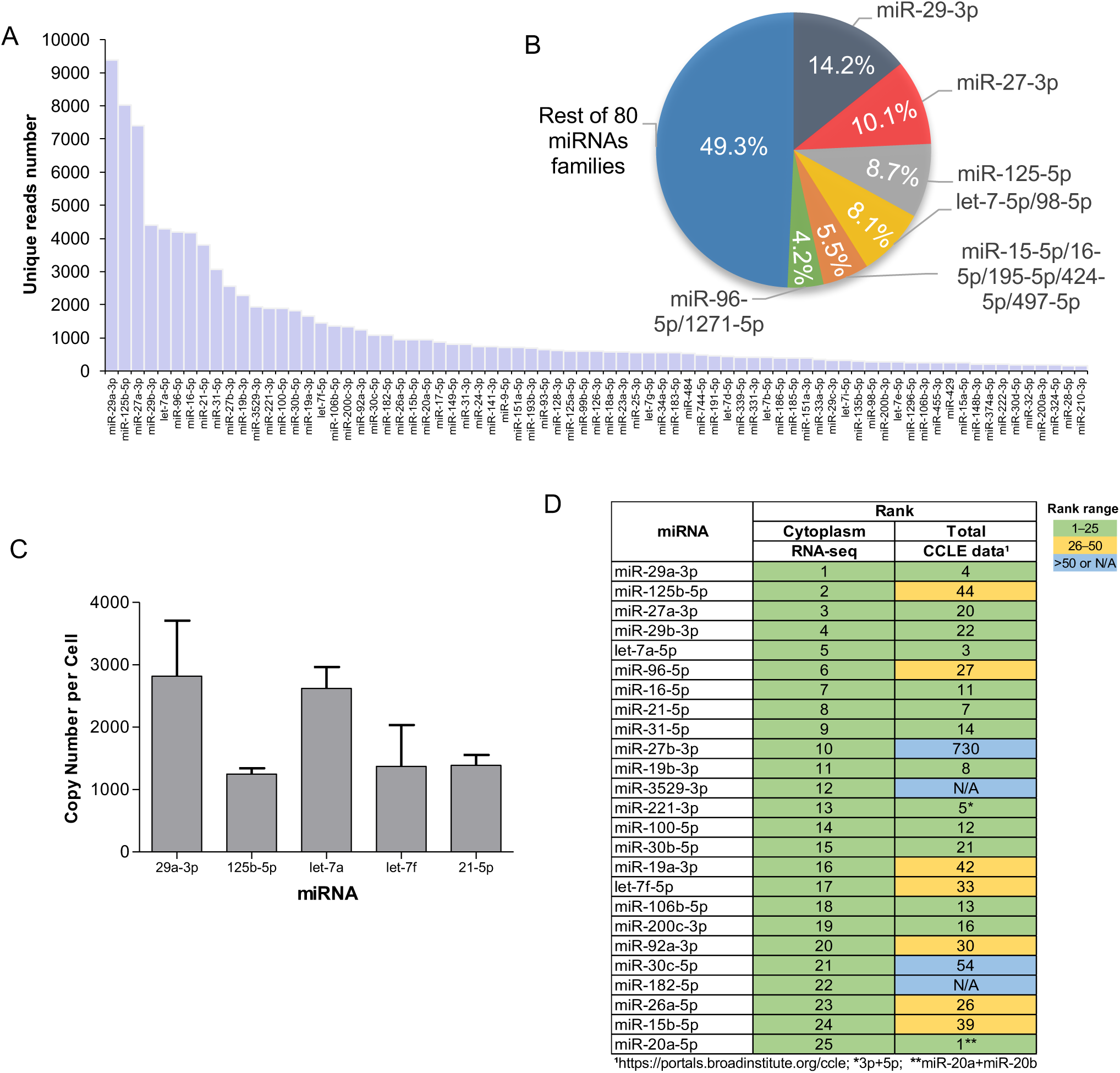
Comparison of miRNAs in HCT116 cells from eCLIP and RIP. (**A**) miRNA abundance in cytoplasm. (**C**) Top 25 most prevalent miRNAs in cytoplasm detected by AGO2-eCLIP-seq compared to detection of those miRNAs identified by Nanostring sequencing of HCT116 cells in published CCLE encyclopedia (Ghandi et al. 2019).

Recently an atlas of miRNA expression was published that used Nanostring sequencing to quantitate miRNA expression (Ghandi et al., 2019). This atlas contains data on hundreds of cell lines and provides a useful background for referencing HCT116 cells. Relative to the Illumina RNAseq used in our study, Nanostring uses different probes and detection strategies. While not an exact comparison, the Nanostring data provide a useful benchmark for putting our measured miRNA expression in HCT116 cells into context

Despite the differences in technologies used for sequencing, thirteen of the top twenty-five miRNAs were shared by our data and data from the atlas describing miRNAs in HCT116 cells (**Figure 3D**). Another seven of the top miRNAs from the atlas were in the top fifty miRNAs identified in our data. The similarity between our measurements of miRNA expression in HCT116 and measurements in other cell lines support the belief that HCT116 cells can offer general lessons that will apply to other cells.

### Testing for relationship between 3’-UTR cluster and gene expression change

We performed RNAseq to identify changes in gene expression due to AGO1, AGO2, AGO1/2, and AGO1/2/3 knockouts relative to wild-type HCT116 cells (**Figure 4**). Because of the recognized importance of the 3’-UTR, we focused our analysis on genes that possessed significant clusters in that region. Some genes also had clusters within the coding sequence in addition to nearby clusters within the 3’-UTR. These clusters were also included in our analysis.

**Figure 4.**
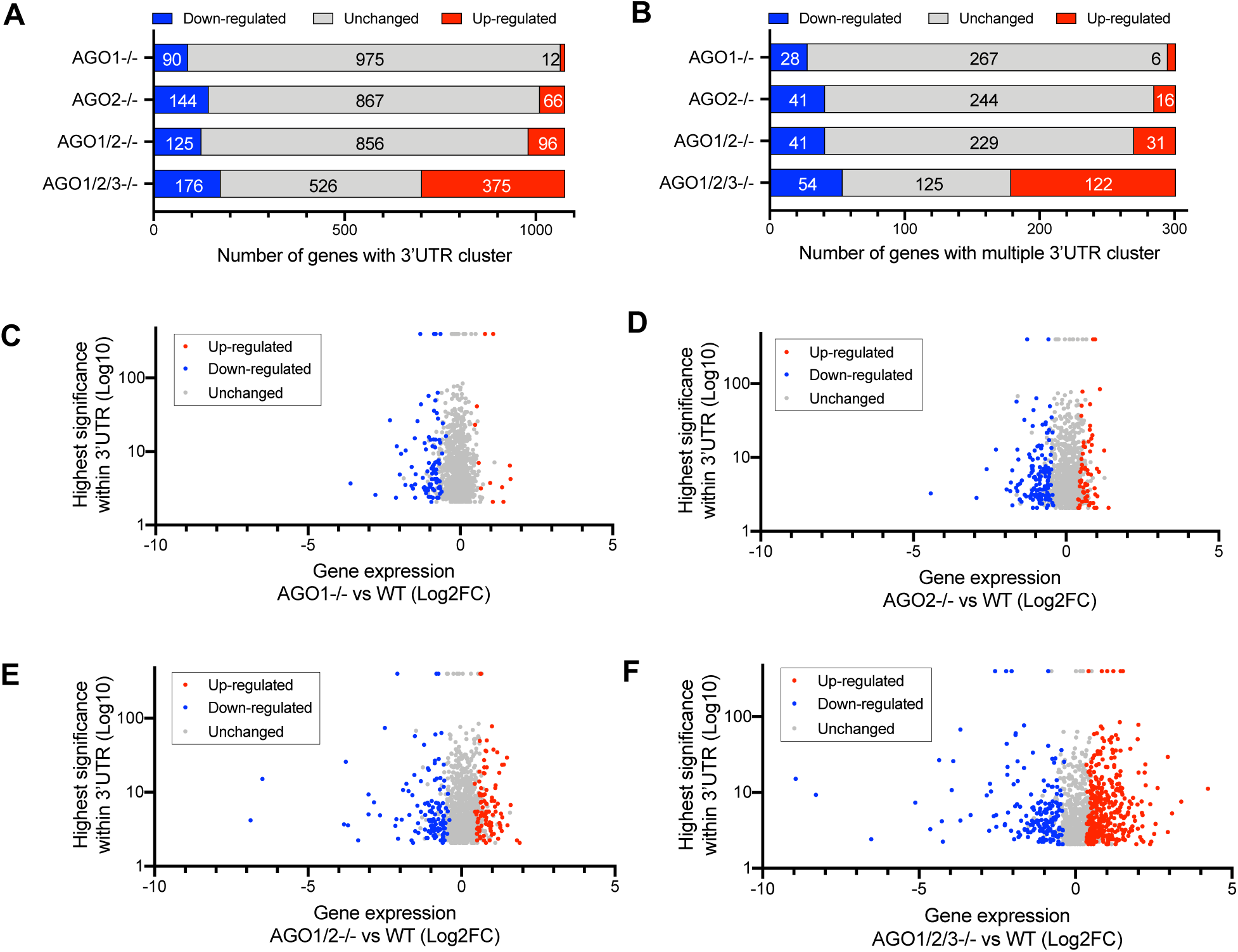
AGO2 binding clusters are associated with up- and down-regulation of gene expression. **(A)** Distribution of gene expression changes for genes with significant AGO2-binding clusters. (B) Distribution of genes with more than one cluster. (**C-F**) Plot of gene expression changes versus cluster significance (**C**) AGO1, (**D**) AGO2, (**E**) AGO1/2, and (**F**) AGO1/2/3 knockout cells.

eCLIP revealed that over 1000 genes possessed AGO2-binding clusters within their 3’-UTRs (**Figure 2**). For AGO1, AGO2, and AGO1/2 knock out cells, contrary to the expectations of canonical miRNA regulation, more genes with significant clusters showed decreased expression than increased expression (**Figure 4A, Supplemental Figure 4**). Most genes with one or more clusters showed no significant change in expression. For the AGO1/2/3 cells, we observed more up-regulated than down-regulated genes. These data showing that the effects of AGO single or double knockouts are less than the triple knockout are consistent with the data on cell proliferation showing the largest decrease in cell growth for triple knockout cells (**Figure 1E**). 301 genes possessed more than one significant cluster and these produced a strikingly similar distribution of gene expression outcomes relative to the entire pool of genes with one or more clusters (**Figure 4B**).

Not only was cluster significance or cluster number not broadly correlated with up- or down-regulation, we also observed little correlation between the absolute values for gene expression change and cluster significance (**Supplemental Figure S4**). Correlation plots showed no significant correlation for the AGO1, AGO2, and AGO1/2 knockout cells for either gene activation or gene repression. For AGO1/2/3 knockout cells there was a slight positive correlation, R=0.19 (p<0.05) for genes with reduced expression and R=0.14 (p<0.05) for genes showing enhanced expression.

### Individual clusters do not predict gene expression change

We individually examined clusters, ranking them by read number and statistical significance relative to clusters in AGO2 knockout cells (**examples shown in Figure 5A-C**). Clusters within 3’-UTRs varied dramatically. Some genes had only one or two clusters. Other genes had multiple clusters in close spatial proximity. The spatial proximity of multiple clusters is likely explained by the observation that more than one AGO protein can bind to the scaffolding protein TNRC6 (Elkayam et al., 2017; Hicks et al., 2017). TNRC6 bridges adjacent AGO:miRNA complexes, leading to cooperative interactions that can strengthen binding (Broderick et al., 2011). Additional clusters suggests the binding of additional AGO:miRNA complexes, and additional complexes would suggest greater binding affinity and potentially great impact on the regulation of gene expression.

**Figure 5.**
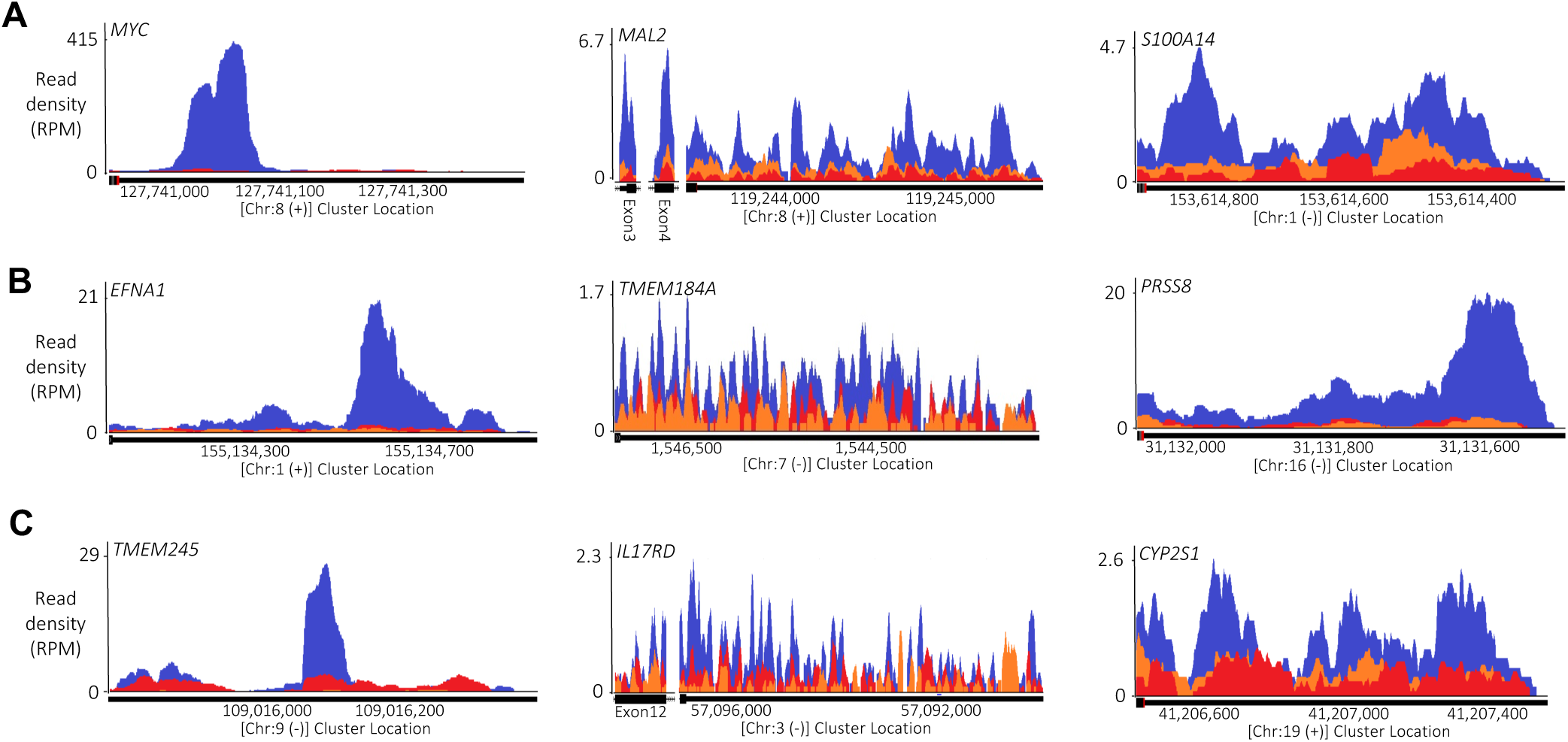
AGO2 binding clusters for genes that are down regulated, do not change, or are up-regulated. (Examples of Ago2 binding clusters identified by Ago2 eCLIP-seq. Blue: Wild type cells. Red: AGO2 knockout cells. Orange: Input control. All 3’-UTR regions, while additional clusters in exons labeled separately. Representative AGO2-binding clusters for (**A**) down-regulated (**B**) unchanged and (**C**) up-regulated genes. See also Figure S3

As we had observed for the overall correlation of cluster significance versus gene expression change, (**Figure 4B**), we observed little correlation between the appearance of individual clusters and gene repression, de-repression, or the absence of significant change (**Figure 5A-C**). For all three categories, some genes were characterized by a single outstanding gene cluster, while others were characterized by multiple clusters.

### Relationship between gene expression changes in knockout cells

In AGO1/2/3 knockout cells, we observe expression changes in 551 genes that contain significant AGO2 read clusters (**Figure 6**). 375 were up-regulated as would have been predicted by the canonical model, while 176 were down-regulated. For genes that are up-regulated upon knockout of one or more AGO variants, the expression of just seven genes is changed in both the AGO1 and AGO1/2/3 knock out cell lines (**Figure 6A**). For genes that are up-regulated upon loss of AGO2, just 13 genes are shared. These results suggest that, by themselves, AGO1 or AGO2 have little impact on repression at genes with 3’-UTR AGO-binding clusters. A much larger number, the expression of 91 genes, was changed in both AGO1/2 and AGO1/2/3 knockout cells.

**Figure 6.**
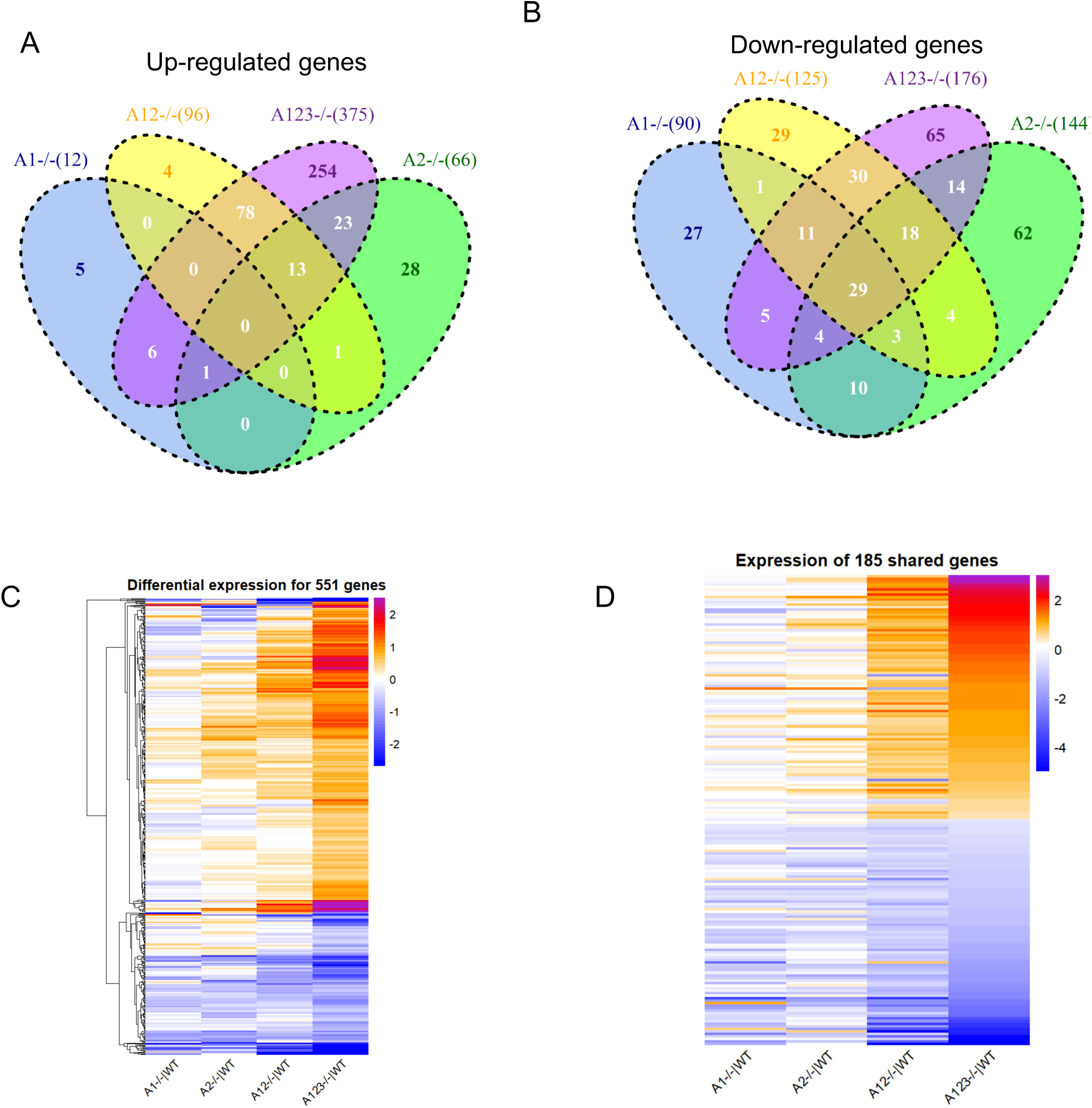
Expression changes in AGO1, AGO2, AGO1/2, and AGO1/2/3 KO cells for genes with AGO binding clusters within their 3’-UTRs. (**A,B**) VENN diagram showing the the overlap among genes with AGO2-binding clusters and significant expression changes relative to wild-type cells for (**A**) increased or (**B**) decreased gene expression after knockouts. (**C**) Heat map showing how the expression relative to wild-type cells changes in AGO1, AGO2, AGO1/2, and AGO1/2/3 cells for the 551 genes associated with AGO2 that are differentially expressed in AGO1/2/3 cells. (**D**) Heat map analysis showing relative changes in gene expression of the 165 AGO-associated genes that show significant expression changes in both AGO1/2 and AGO1/2/3 knockout cells.

For genes that are down-regulated upon knockout of one or more AGO variants, more genes were shared between the AGO1 or AGO2 single knockout cells and AGO1/2/3 triple knockout cells than had been the case for up-regulated genes (**Figure 6B**). 49 genes were shared between the AGO1 and AGO1/2/3 knockout cells, while 51 were shared between AGO2 and AGO1/2/3 knock out cells. These data sugges that AGO1 and AGO2 may have more impact on increasing gene expression than on decreasing gene expression. The expression 88 genes were changes in both AGO1/2 and AGO1/2/3 cells.

We constructed a heatmap to visualize the relationship between the changes in gene expression for the 551 AGO2-associated genes with altered expression in the AGO1/2/3 knockout cells (**Figure 6C**). The magnitude of change is relatively small in AGO1 or AGO knockout cells, becomes larger in the AGO1/2 knockout cells, and reaches a maximum in AGO1/2/3 knockout cells. Visual inspection suggests that the trends in rank order for expression changes are similar in the four cell lines.

The AGO1/2/3 knockout cells growth more slowly that the other knockout cell lines, raising the possibility that there are more widespread disruptions in gene expression that are not directly related to AGO expression or the potential for any interactions with mRNA. Therefore, we focused on the 185 genes that are altered in both AGO1/2 and AGO1/2/3 knockout cells as having a high potential to be more directly related to a knockout of AGO protein rather than a more indirect effect from global gene changes that are reflected in slower cell growth.

Comparison of AGO1/2 and AGO1/2/3 data revealed a strong correlation between the gene expression changes observed for individual genes in the two cell lines. Heatmap representation of these gene expression changes revealed a strong similarity among the four AGO knockout cell lines for gene repression and depression (**Figure 6D**). There is a strong trend among all four AGO variants separating down-regulated and up-regulated genes. The magnitude of change reached the greatest magnitude in the AGO1/2/3 knockout cells. These data reinforce the conclusion that AGO proteins exert similar effects and that those effects become more pronounced as the combined level of AGO protein is reduced.

### Analysis of individual clusters within 3’-untranslated regions

We chose 22 genes for detailed expression analysis and comparison. Genes were chosen: 1) based on possession of at least one highly significant read cluster within the 3’-UTR and 2) to represent gene targets with both one and multiple AGO2-binding clusters (**Figure 7A, Supplementary Figure S7A-C)**. Three genes (IL17RD, TMEM245, MAL2) also had one or more AGO binding clusters within the coding region near the junction between the coding region and the 3’-UTR. While the potential for miRNA complementarity was not a criterion for choosing these genes, eighteen of the twenty-two had at least one calculated site of seed-sequence complementarity for a miRNA ranked in the top 25 for prevalence by our RNAseq (**Figure 3, Supplementary Figure S7D**). Several had multiple sites for seed sequence complementarity with highly ranked miRNAs.

**Figure 7.**
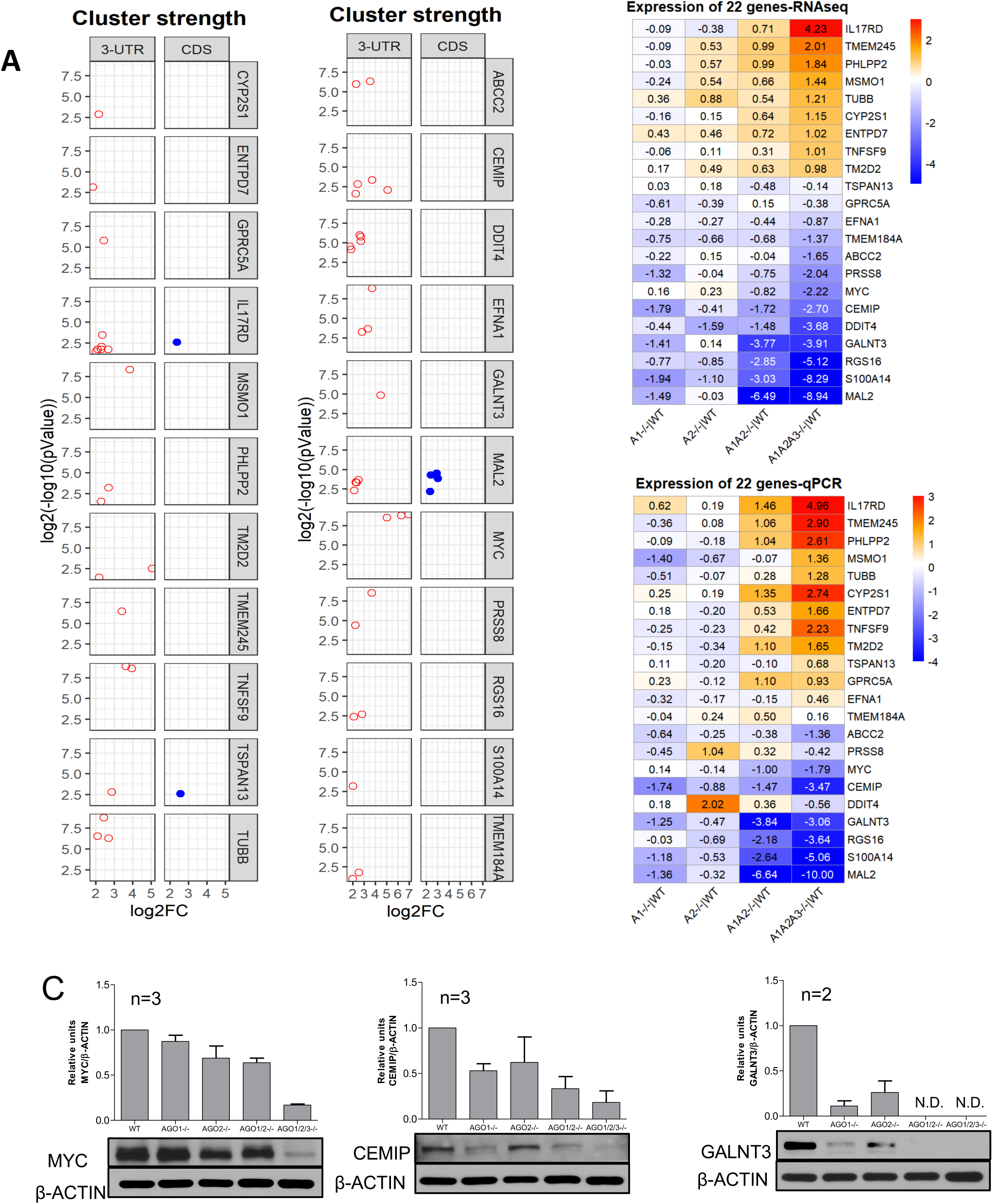
Impact of 22 representative AGO2-binding clusters on gene expression. (**A**) Cluster strength, significance, and location. (**A**) Expression level relative to wild type for 22 selected genes, evaluated by RNAseq and ranked highest to lowest. (**B**) Expression level relative to wild type for 22 selected genes, evaluated by qPCR. Ordering is based on RNAseq rankings. (**C**) Western analysis of MYC, CEMIP, and GALNT3 in wild type and knockout.

As we had observed for the entire group of AGO-associated genes, among the twenty-two genes RNAseq data revealed both gene repression, de-repression, and the absence of significance change (**Figure 7B, Supplemental Figure S7A-C, E**). The changes in the AGO1 or AGO2 knockout cells are relatively muted. For AGO1 knockout cells, five genes show a 2-fold (>1 log_2_-fold) reduction in gene expression and no genes show a 2-fold (>0.5 log2-fold) increase. For AGO2 knockout cells, two genes showed a > 2-fold reduction in expression while four genes show a >2-fold decrease in expression.

The changes in AGO1/2 and AGO1/2/3 knockout cells are greater and there is more similarity between the cell lines. Six genes that show >2-fold reduction in gene expression in AGO1/2 knockout cells and eight genes showed a >2-fold increase. Most of these genes were among the 20/22 genes showing a >2-fold increase or decrease in AGO1/2/3 cells. The rank order for most genes relative to one another was similar. Genes that were more repressed or activated in AGO1/2 knockout cells tended to also be among the most repressed or activated genes in AGO1/2/3 knockout cells. The trend for the magnitude of gene expression to be most altered in AGO1/2 and AGO1/2/3 knockout cells relatively to the AGO1 or AGO2 single knockouts is consistent with the larger number of genes that show significantly altered expression (**Figures 4 and 5**).

We crosschecked the RNAseq data for the 22 genes using quantitative PCR (qPCR) (**Figure 7B, Supplemental Table S1**). For AGO 1 knockout cells, four of the five genes with >2-fold altered expression in our RNAseq data also showed >2-fold change when assayed by qPCR. The AGO2 knockout cells that had shown the least change by RNAseq also showed the least change by qPCR. The qPCR data for AGO1/2 and AGO1/2/3 knockout again showed the greatest changes, with >2-fold changes in expression observed for thirteen and eighteen genes respectively. We analyzed protein expression for three genes that had antibodies that were adequate for western analysis. We observe decreases in MYC, CEMIP, and GALNT3 protein expression as predicted by RNAseq and qPCR (**Figure 7C, Supplemental Figure S7**).

## DISCUSSION

### Reevaluating the standard model linking AGO binding to gene repression

The premise underlying the standard model for the action of miRNAs is that miRNA:AGO complexes recognize sequences within the 3’-UTR. This recognition is primarily through “seed sequence” pairing of bases 2-7 at the 5’ end of miRNAs, with the potential for other bases to contribute (Lewis et al., 2003; Grimson et al., 2007). Recognition is associated with miRNA-mediated translational repression of these genes, with hundreds or thousands of genes predicted to be subject to this repressive mechanism (Friedman et al., 2009). Repression is often inferred based on the seed sequence complementarity, helping shape the hypotheses of thousands of publications each year. In accord with these common assumptions, AGO binding detected by CLIP within a 3’-UTR should be a major factor predicting gene repression by miRNAs. Our goal was to test this hypothesis by combining AGO knockout cells, RNAseq, and anti-AGO2 eCLIP (**Figure 1**).

### Requirement for knocking out AGO1, AGO2, and AGO3

We evaluated gene expression in AGO1, AGO2, AGO1/2, and AGO1/2/3 knockout cells. Trends for gene expression are similar in all knockout cell lines (**Figure 5B,C**), suggesting that the AGO variants perform similar functions. Absolute levels of gene expression change vary dramatically. Individual knockouts of AGO1 or AGO2 have much less impact on overall gene expression than AGO1/2 double knockout while the AGO1/2/3 triple knockout has the largest impact on gene expression. Consistent with these data, cell growth of the single knockout cells is closer to wild-type than the double knockout cells, with cell growth of the AGO1/2/3 knockout cells being the slowest.

While our data do not exclude the possibility that the AGO variants have unique functions, our data suggest that AGO1, AGO2, and AGO3 have redundant functions that tend to produce similar changes in gene expression. Overlapping functions for AGO variants has previously been suggested (Su et al. 2009; Ruda et al., 2014). It had previously been shown that FLAG-HA-tagged AGO1-4 bind to similar sites within the transcriptome (Hafner et al., 2010) supporting the conclusion that endogenously expressed AGO1 and AGO3 will also bind to sites like the ones where we observe AGO2 association.

### AGO-binding Clusters are associated with gene repression and de-repression

It would have been unrealistic to expect that knockdown of AGO variants would only result in de-repression of genes (Gosline et al., 2016). A change in the expression of one gene will inevitably produce changes in the expression of other genes that are unrelated to association of AGO with those mRNAs. These indirect secondary effects on gene expression would be expected to yield confounding effects that would confuse analysis. Nevertheless, based on standard assumptions, we had assumed that the strength and significance of AGO binding would be an important factor predicting gene de-repression upon knocking out AGO expression.

Our data indicate do not reveal a predictive link between AGO binding and gene repression (**Figures 4-7**). Some genes with AGO binding clusters within their 3’-UTRs showed the expected increased expression – the expected outcome. However, for AGO1, AGO2, and AGO1/2 knockout cells, more genes showed decreased expression. For AGO1/2/3 knockout cells, more genes showed increased expression as might have been predicted by the standard model, but expression for a substantial number decreased. Not only did we observe both increases and decreases in gene expression, we also observed that most genes with AGO-binding clusters did not exhibit changes in expression.

There was no predictive correlation between the significance or magnitude of AGO-binding clusters and gene repression or de-repression. Inspection of hundreds of gene clusters individually revealed that clusters associated with gene repression, activation, or lack of significant change were indistinguishable.

### *MYC*: Strong cluster associated with AGO-induced gene activation

*MYC* is an oncogene whose overexpression can lead to cancer (Dang, 2012). Several previous studies have concluded that *MYC* expression can be down-regulated by miRNAs (Kim et al., 2009; Sampson et al., 2007; Swier et al., 2019). Our eCLIP data showed that the strongest read cluster within any 3’-UTR in our dataset was located within the *MYC* 3’-UTR. Let-7 is one of the most highly expressed RNAs in HCT116 cells, and there is a strong Let-7 site within the *MYC* 3’-UTR (**Supplemental Figure S7D and 8**). The simplest conclusion from combining our eCLIP data with standard assumptions about miRNAs, the existence of a likely Let-7 target site, and the previous reports of miRNA action would be that *MYC* expression is repressed by miRNAs in HCT116 cells. This conclusion would then lead to the hypothesis that the repression might be important for understanding the regulation of *MYC* expression during cancer.

However, when we correlated the association of AGO2 with difference in expression of *MYC* for wild-type versus AGO1/2/3 knock out cells, we observed the opposite result. In AGO knockout cells, the expression of *MYC* RNA and protein decreases relative to wild-type cells. Rather than repress *Myc* expression, our data suggest that the RNAi machinery promotes expression.

Whether this regulation of *MYC* expression is direct or indirect is not clear and conclusively determining the mechanism is beyond the scope of this study. Increased expression might be due to a non-canonical mechanism in which a miRNA binds the 3’-UTR and activates gene expression. *MYC* is a highly expressed gene, it is possible that it might act as a miRNA “sponge” to attract enough miRNAs to affect global regulatory pathways. Alternatively, knocking out AGO may trigger changes in gene expression that indirectly enhance *MYC* expression.

Regardless of which mechanism is responsible for reduced MYC expression, the simple assumption from our eCLIP data combined with prior dogma, that *MYC* expression is repressed by miRNA:AGO complex, is incorrect. *MYC* is a critical gene for understanding cancer. Making progress towards understanding the potential for miRNAs to control *MYC* and other genes underlying disease will benefit from reexamining prior assumptions.

## Conclusions

The control of mammalian gene expression by miRNAs has been the subject of intense interest for almost twenty years. While the interplay of RNA and gene expression is known to be a complex phenomenon (Cech and Steitz, 2014), many reports have revolved around the simple assumption that recognition by AGO:miRNA complexes leads to repression of gene expression. Contrary to these assumptions, our data suggest that AGO-bound sequences within 3’-UTRs are not reliable predictors of gene repression. If eCLIP data do not correlate well with the potential for miRNA-mediated gene repression, studies that rely only on computationally predicted seed matches for miRNAs will be even more problematic.

Assumptions about the mechanism of action of miRNAs in their endogenous context should be carefully justified. Gene repression based on seed sequence matches should not be assumed. Our findings raise questions about the origins and mechanistic significance of AGO-bound clusters that are not associated with gene repression. Answering these questions, and the further questions that will arise from address them, will be required to put the understanding of miRNA-mediated regulation and improve the likelihood of important insights into basic science and therapeutic development.

Our data do not dispute the canonical mechanism that AGO:miRNA complexes can bind to 3’-UTRs and repress gene expression. That conclusion has been amply proven by many studies using 3’-UTRs engineered to include binding sites for miRNAs (Bartel, 2018). Our data do suggest that inferring an association between repression of endogenous gene expression and binding of miRNA:AGO complexes at a site is not straightforward. It is useful to consider that the standard mechanism should not be viewed as the only possible mechanism for the action of miRNAs. While activation of genes like *MYC* effects may be indirect, they may also be through a novel mechanism. Researchers should keep an open mind to the fact that AGO may regulate the action of target RNA through non-canonical mechanisms beyond repression of translation in cell cytoplasm.

## SUPPLEMENTARY DATA

## ACKNOWLEDGEMENTS

The authors thank Dr. Joshua Mendell for the gift of HCT116:AGO2 knock out cells used in eCLIP experiments.

## FIGURES

**Figure Supplementary Figure 1A (related to Figure 1).**
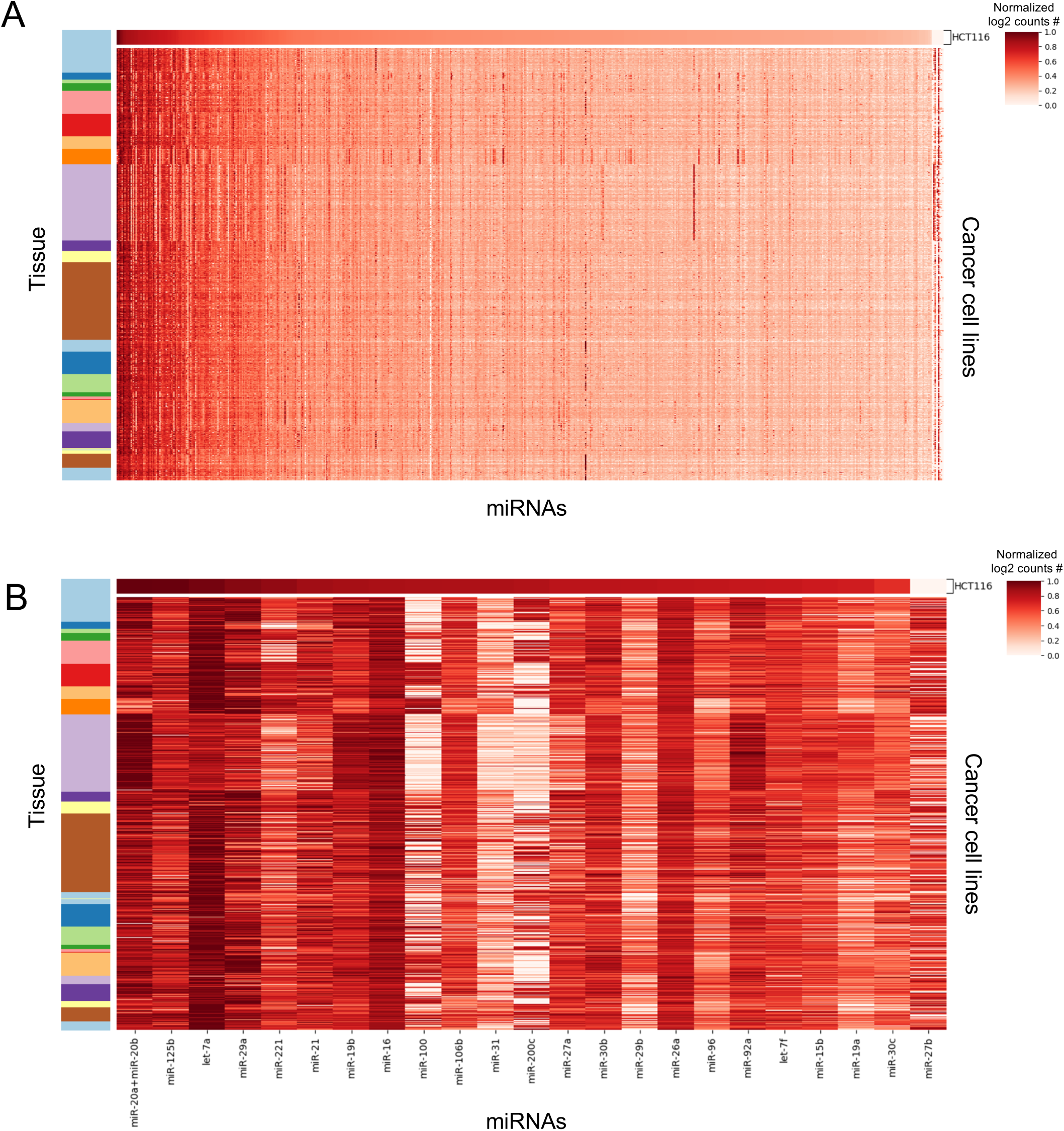
miRNA expression profile in cancel cells based on Cancer Cell Line Encyclopedia (CCLE). (**A**) Relative expression or miRNAs cancer cell lines. (**B**) Relative expression of the most highly expressed miRNAs in HCT116 cells and other cell lines. For (**A**) and (**B**), The top lines represent expression of miRNAs in HCT116 cells and show similar trends for miRNA expression from higher expressed to less highly expressed. miRNAs ordered from HCT116 mostly to lowest expressed. Tissue color bar order from top to bottom (LARGE INTESTINE, AUTONOMIC GANGLIA, BILIARY TRACT, BONE, BREAST, CNS, ENDOMETRIUM, FIBROBLAST, HAEMATOPOIETIC AND LYMPHOID, KIDNEY, LIVER, LUNG, OESOPHAGUS, OVARY, PANCREAS, PLEURA, PROSTATE, SALIVARY GLAND, SKIN, SMALL, SOFT TISSUE, STOMACH, THYROID, UPPER AERODIGESTIVE TRACT, URINARY TRACT).

**Figure S2 (Related to Figure 2).**
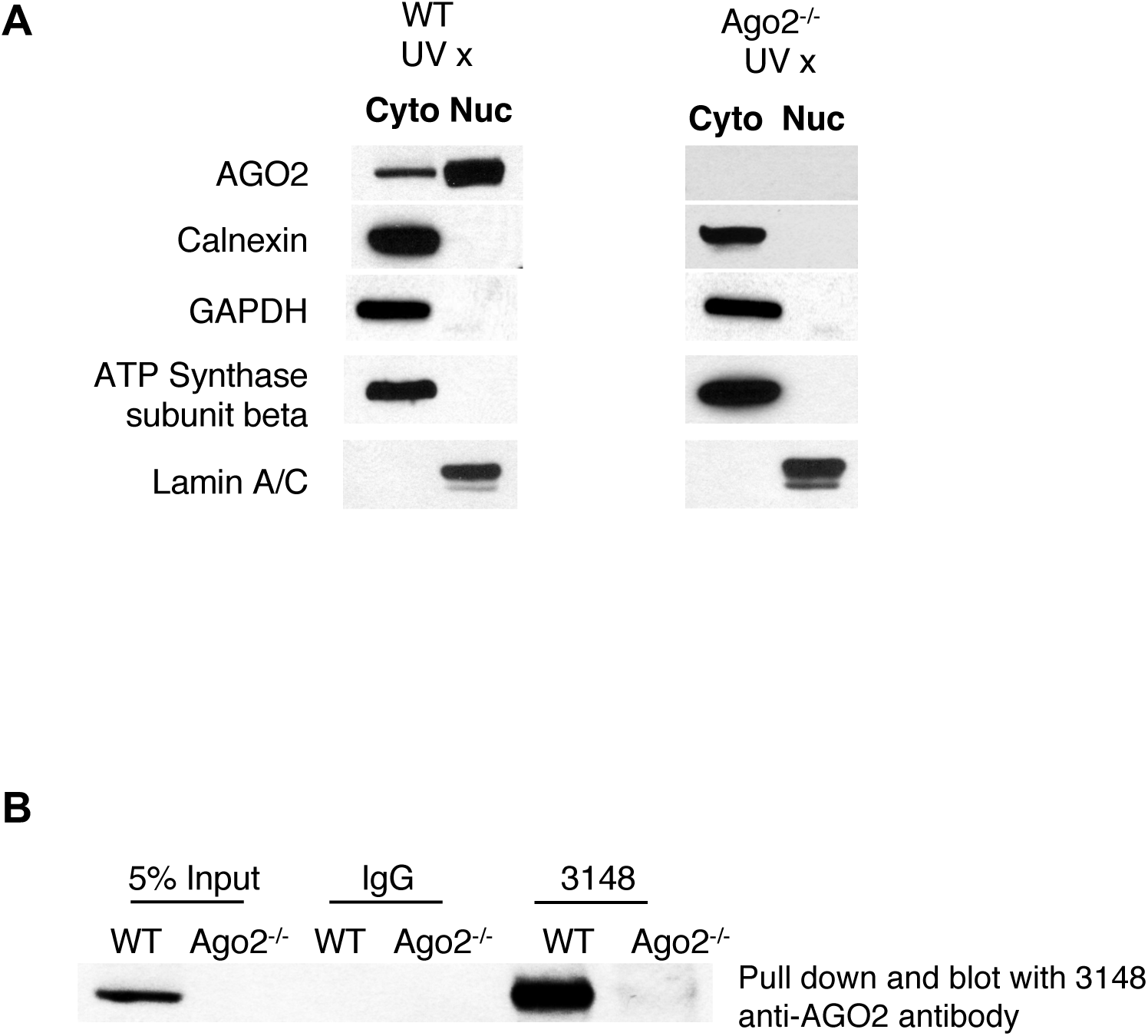
Western blot showing (B) purity of cytoplasm prep for eCLIP and (C) pull down using anti-AGO2 rabbit polyclonal antibody (3148).

**Figure S3 (Related to Figure 2).**
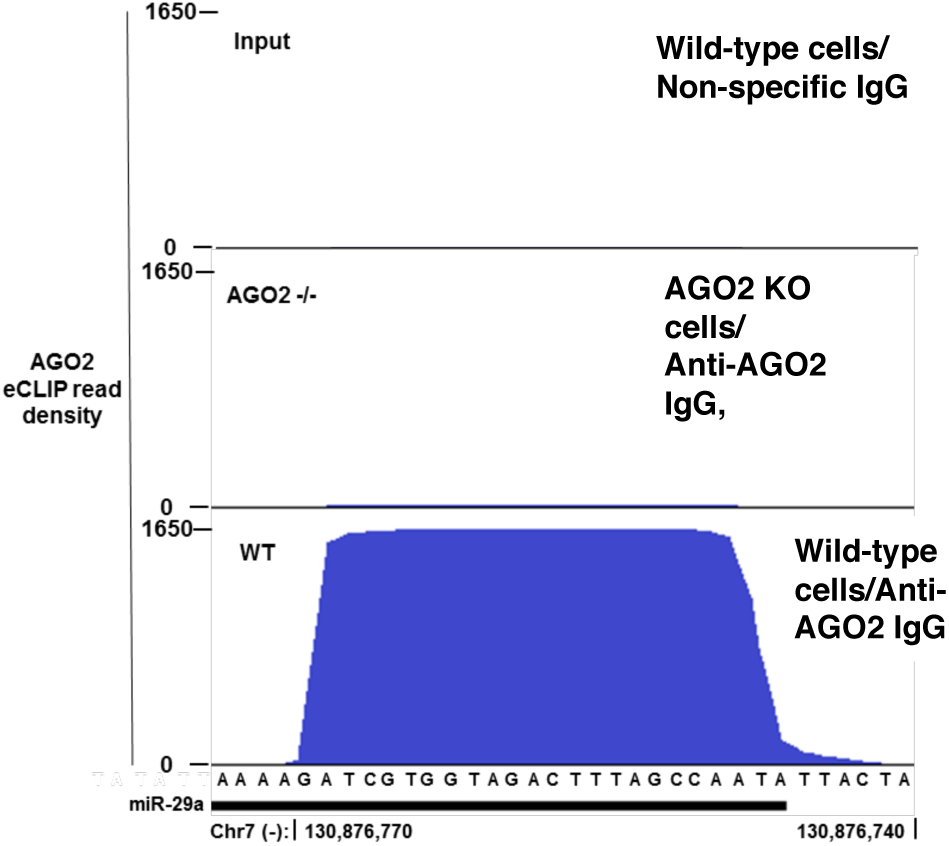
Sample cluster data for highly ranked cluster – miR-29a.

**Supplemental Figure S4 (Related to Figure 4).**
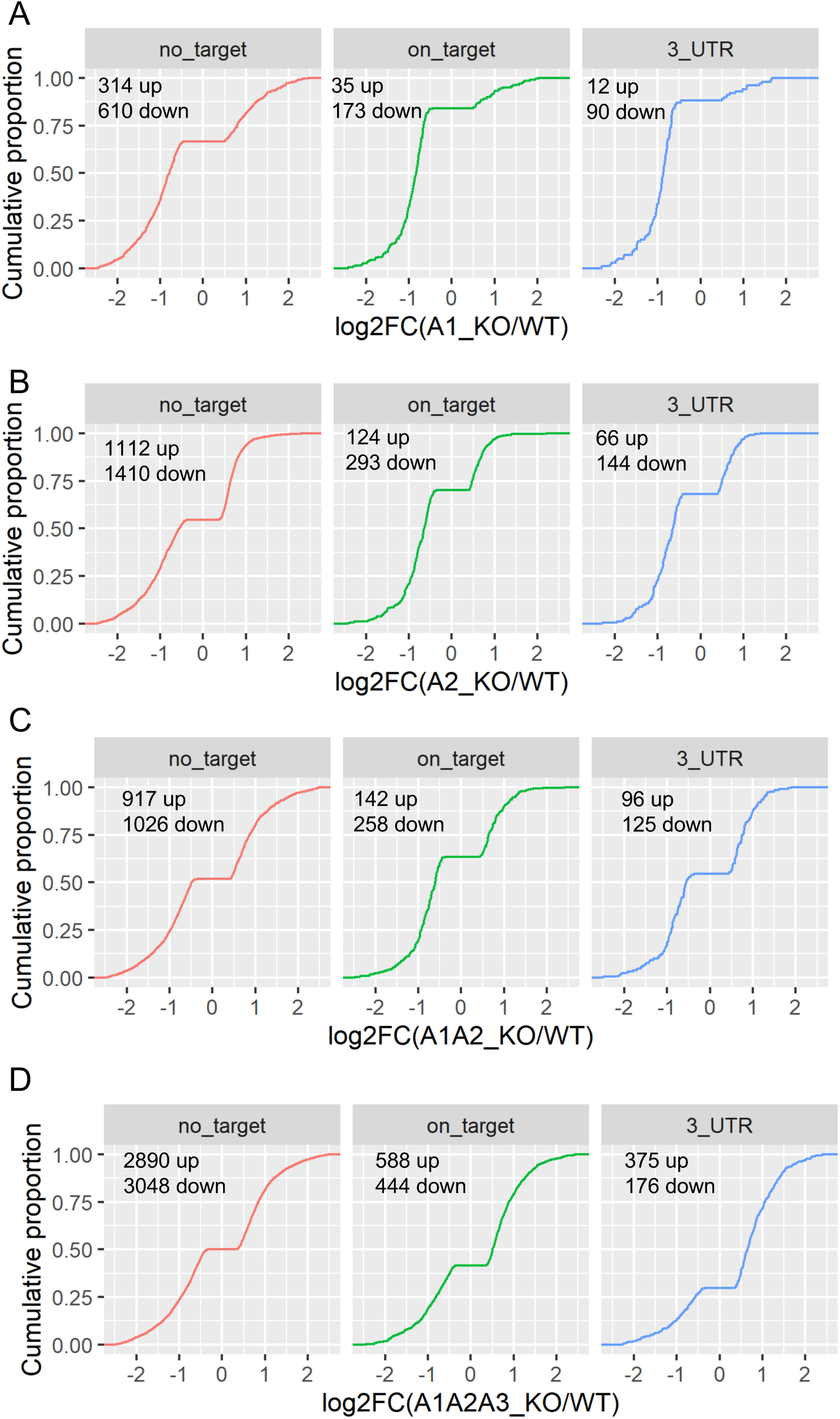
AGOs role in genes regulation in HT116 cells. (**A-C**) CDF plot of log2 fold change upon Ago KO for genes having no Ago2-binding clusters, genes that have AGO2 clusters, and genes that have AGO2 clusters with their 3’-UTRs. (**A-D**) Data for AGO1, AGO2, AGO1/2, and AGO1/2/3 knockout cells respectively.

**Supplemental Figure S4 (Related to Figure 4).**
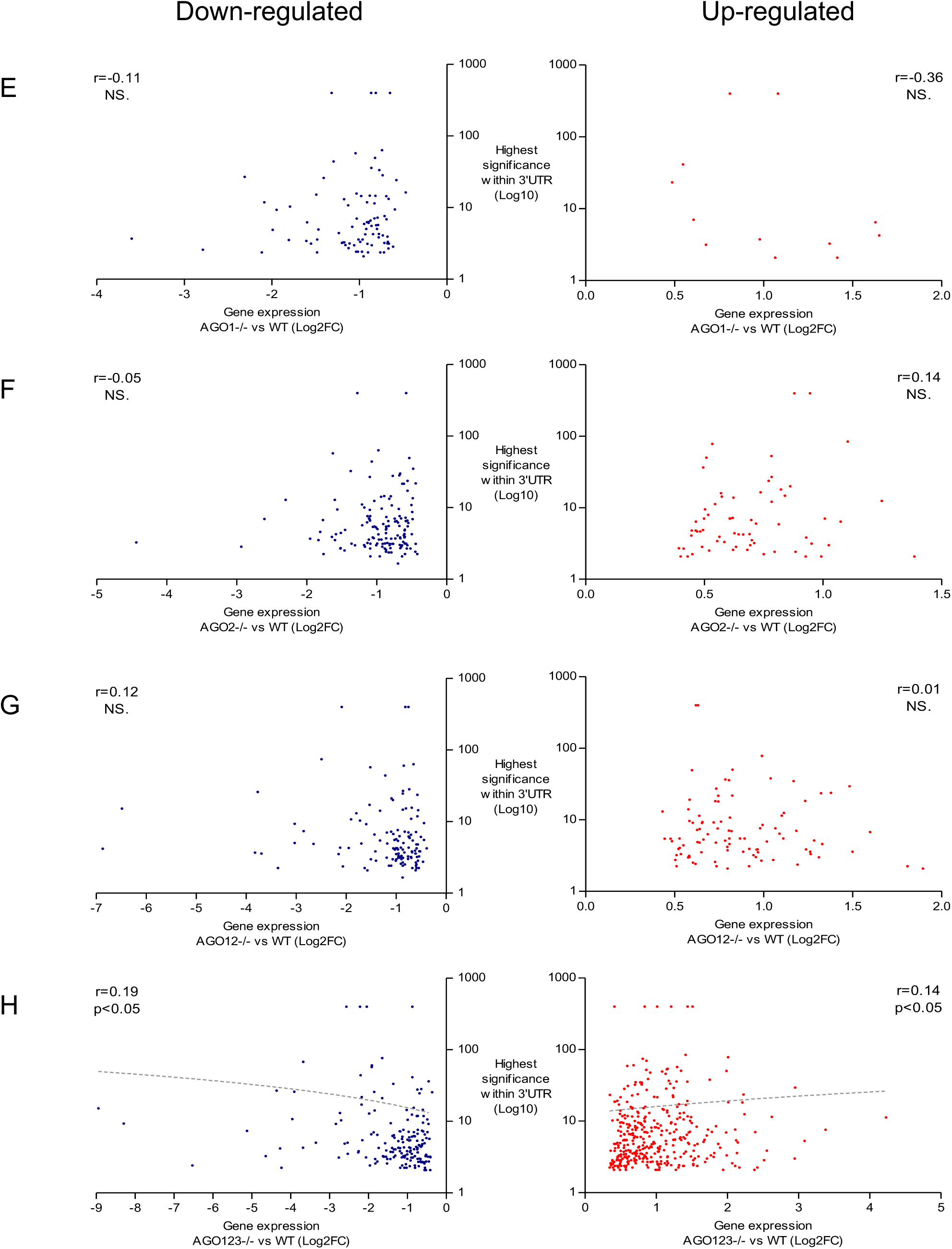
Correlation between significance of AGO2 binding cluster and gene expression change in AGO KO cell lines. **(E-H)** Data for AGO1, AGO2, AGO1/2, and AGO1/2/3 knockout cells respectively. Blue: downregulated genes. Red: Up-regulated genes. Data do not pass normality test for both data sets, or relation was not linear, therefore Spearman correlation was applied.

**Supplemental Figure S3.**
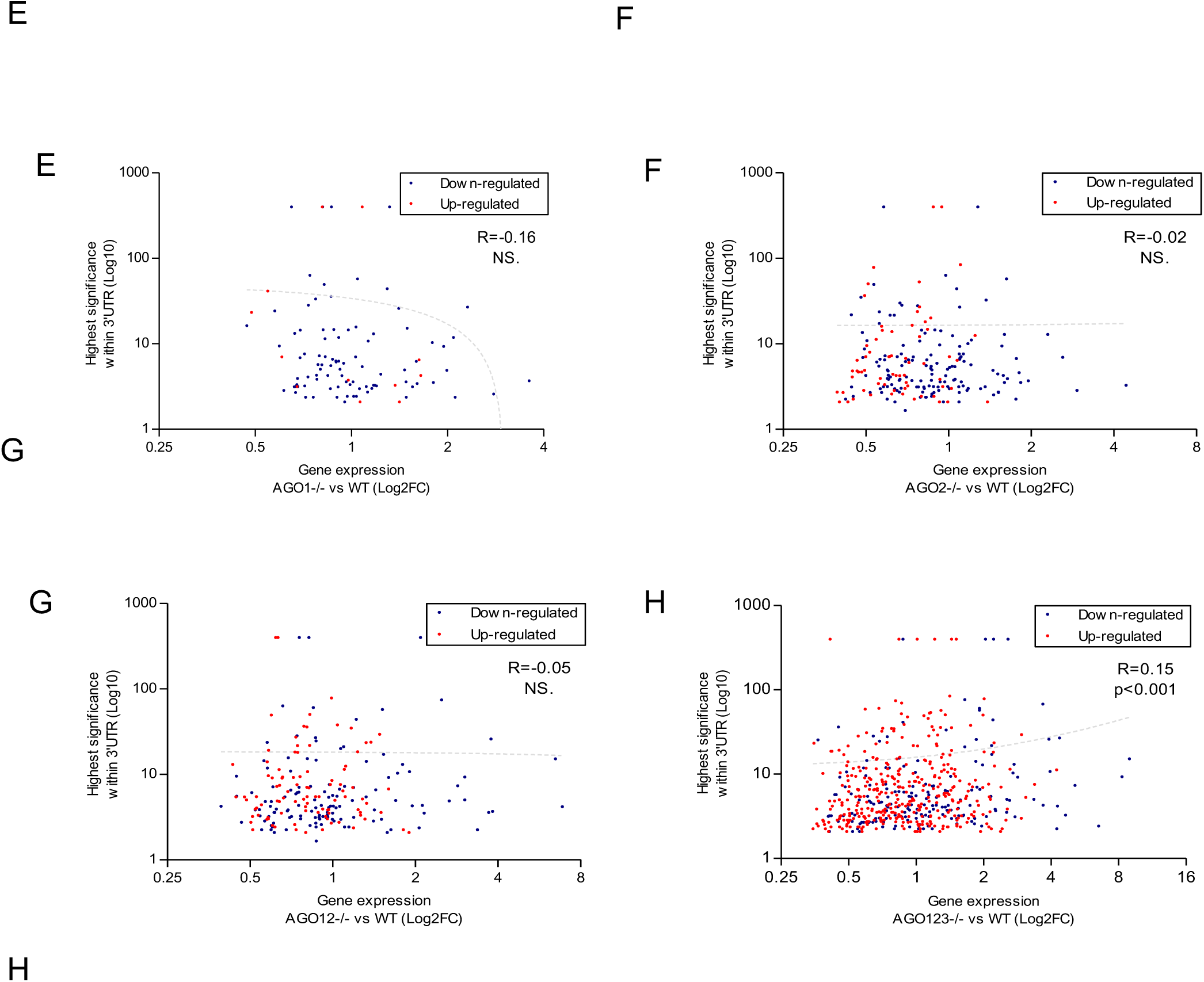
AGOs role in genes regulation in HT116 cells. (**A-C**) CDF plot of log2 fold change upon Ago KO for genes having no Ago2-binding clusters, genes that have AGO2 clusters, and genes that have AGO2 clusters with their 3’-UTRs. (**A-D**) Data for AGO1, AGO2, AGO1/2, and AGO1/2/3 knockout cells respectively. (**E-H**) Plot of gene expression changes versus cluster significance correlation (**E**) AGO1, (**F**) AGO2, (**G**) AGO1/2, and (**H**) AGO1/2/3 knockout cells. Data do not pass normality test, therefore Spearman correlation was applied.

**Supplemental Figure 7a.**
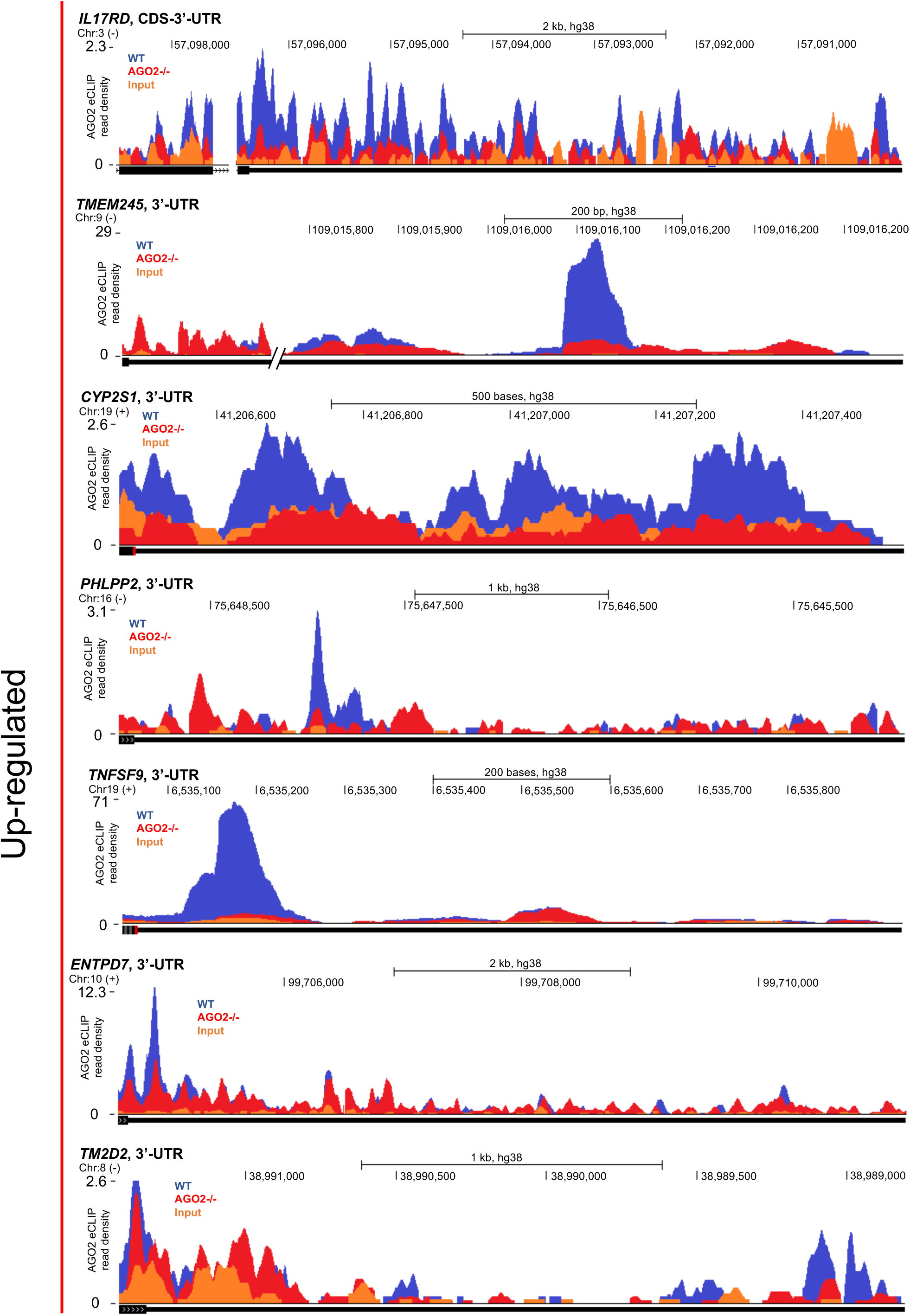

**Supplemental Figure S7B.**
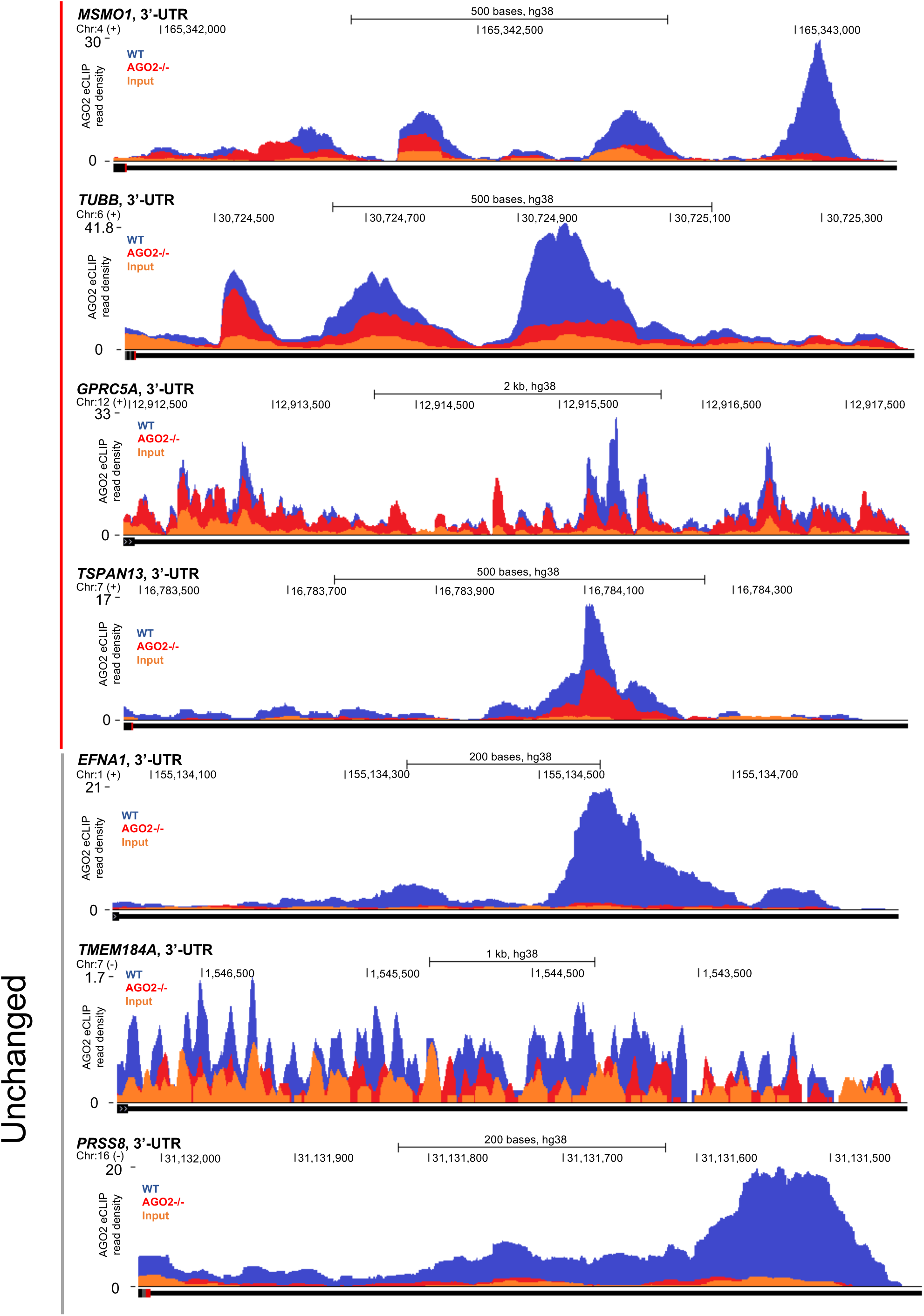

**Supplemental Figure S7C.**
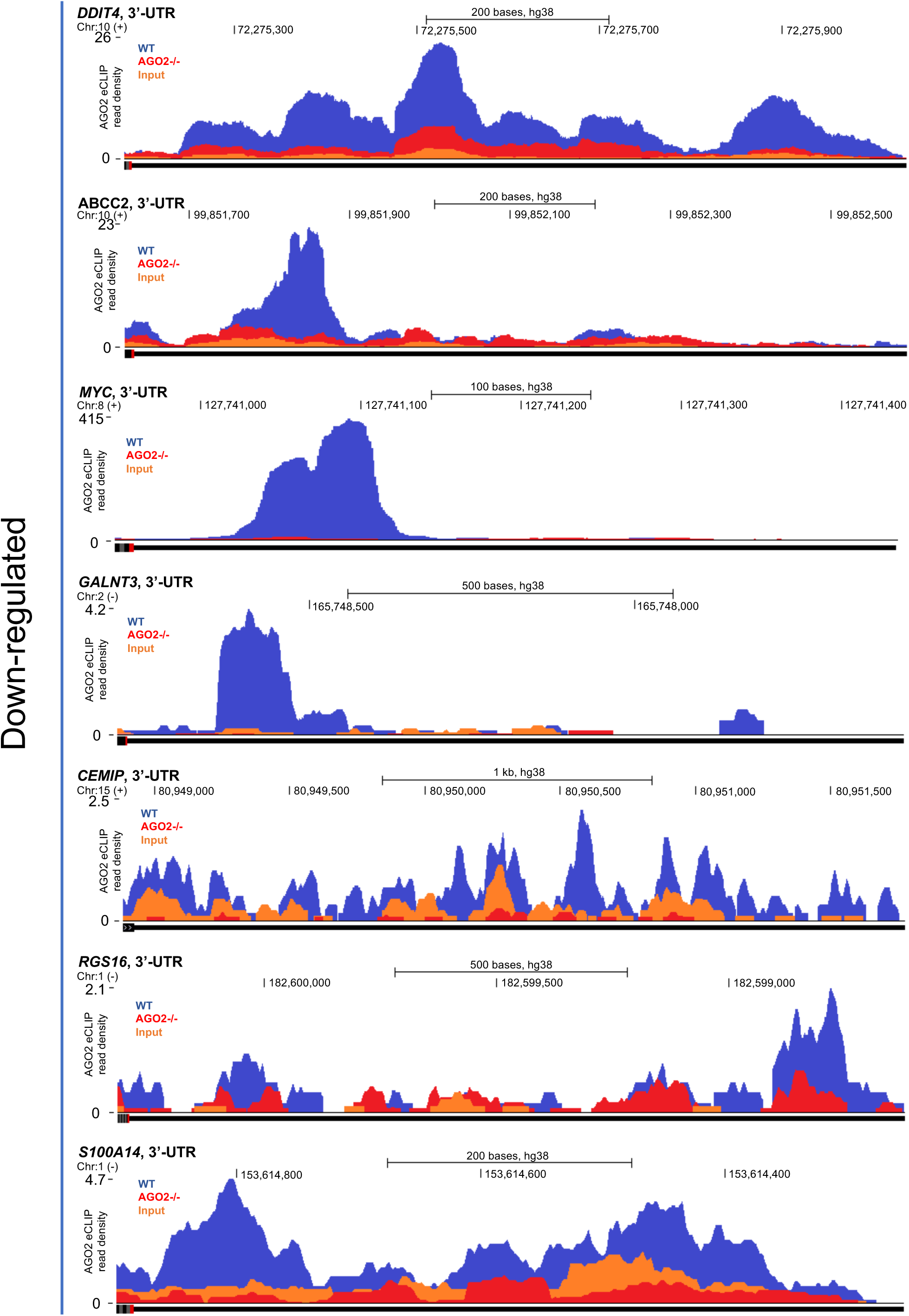
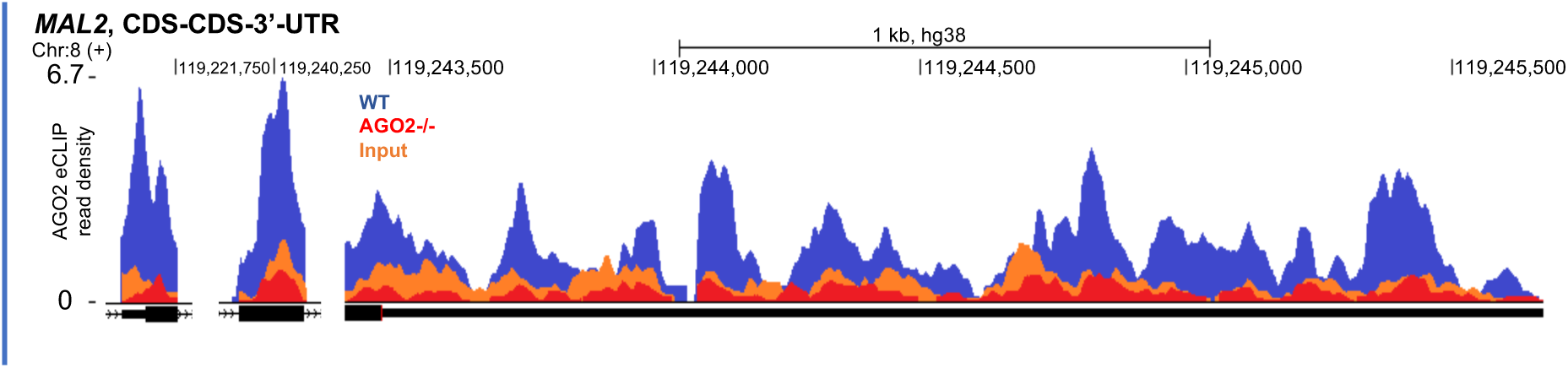

**Supplemental Figure_S7D.**
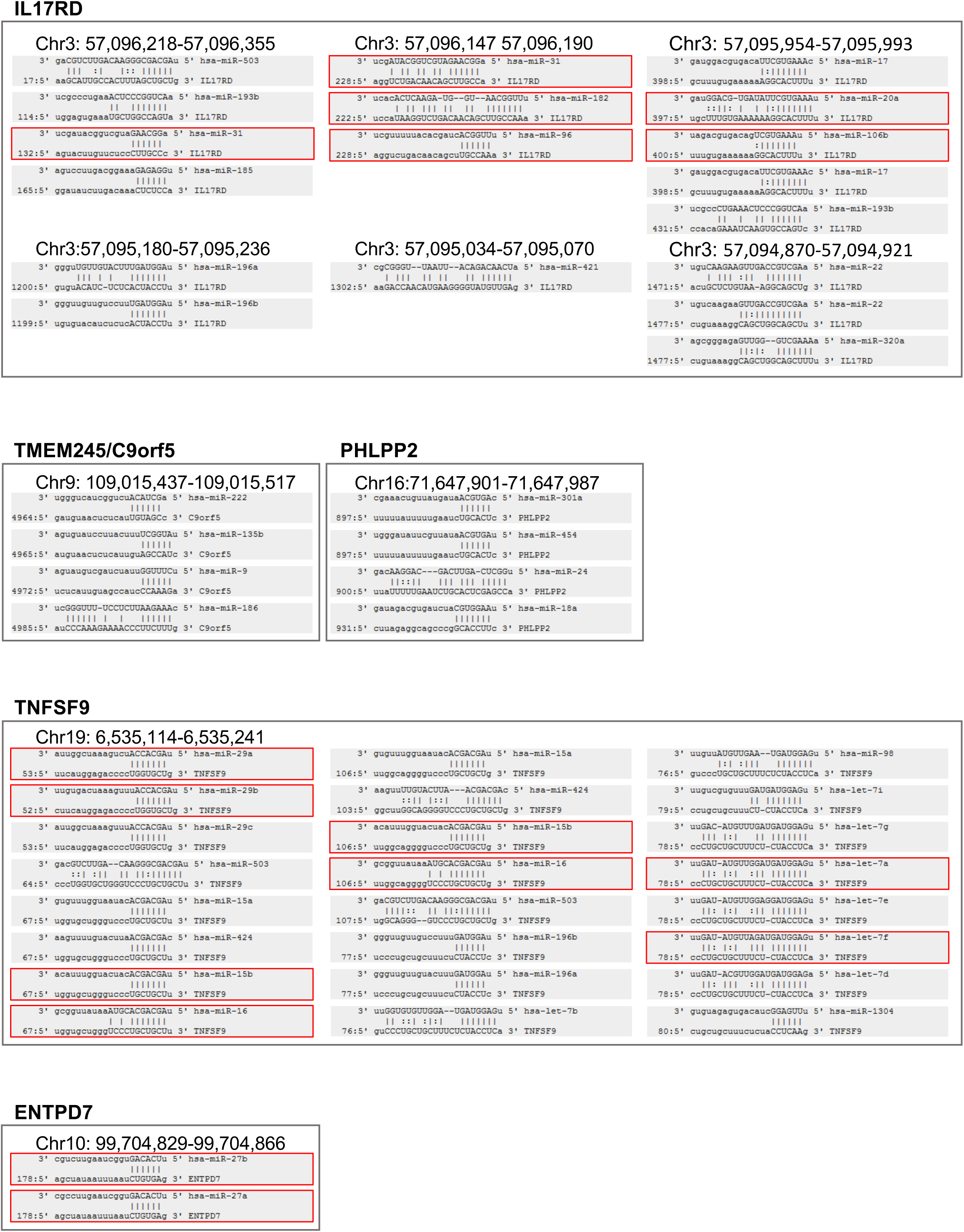
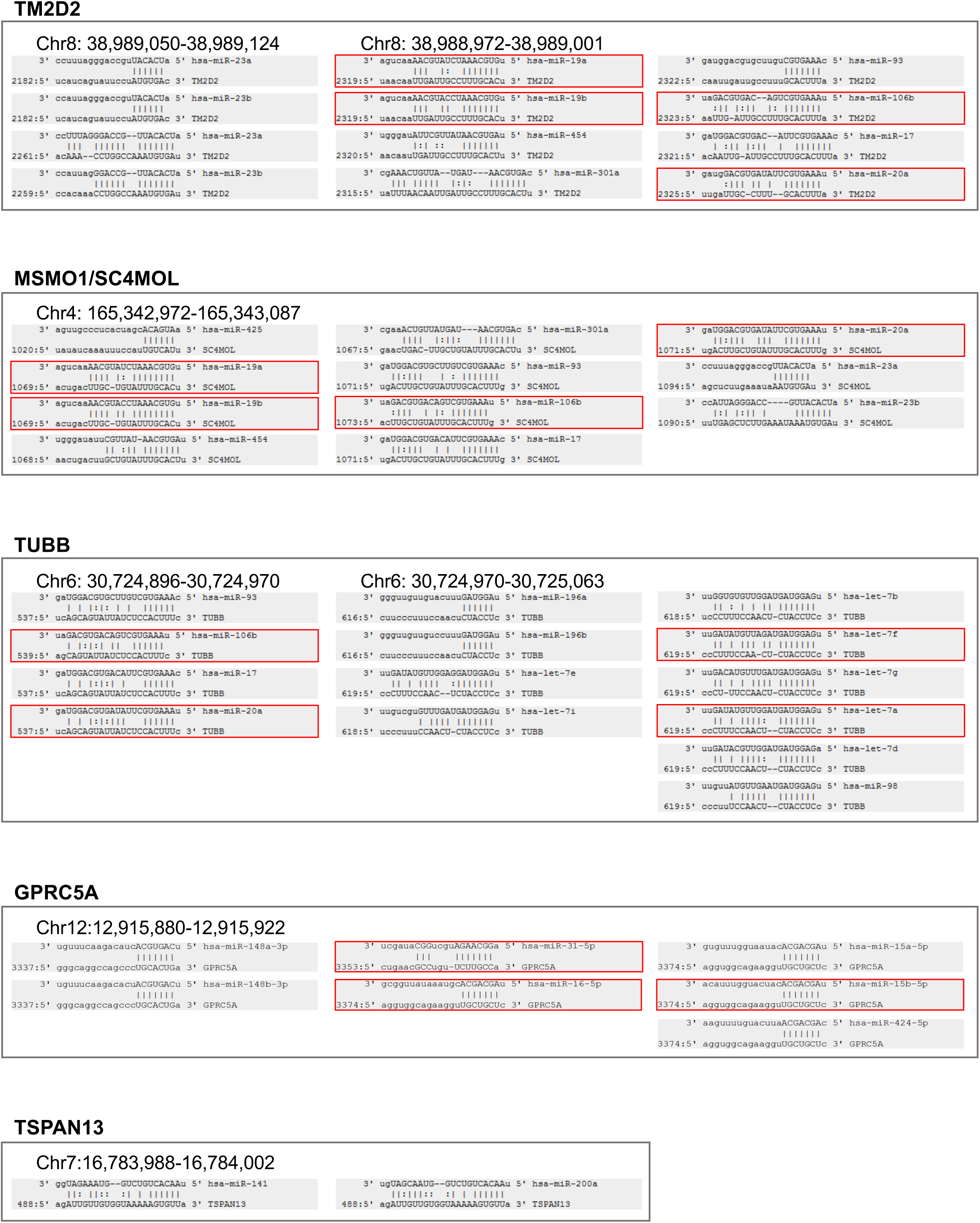
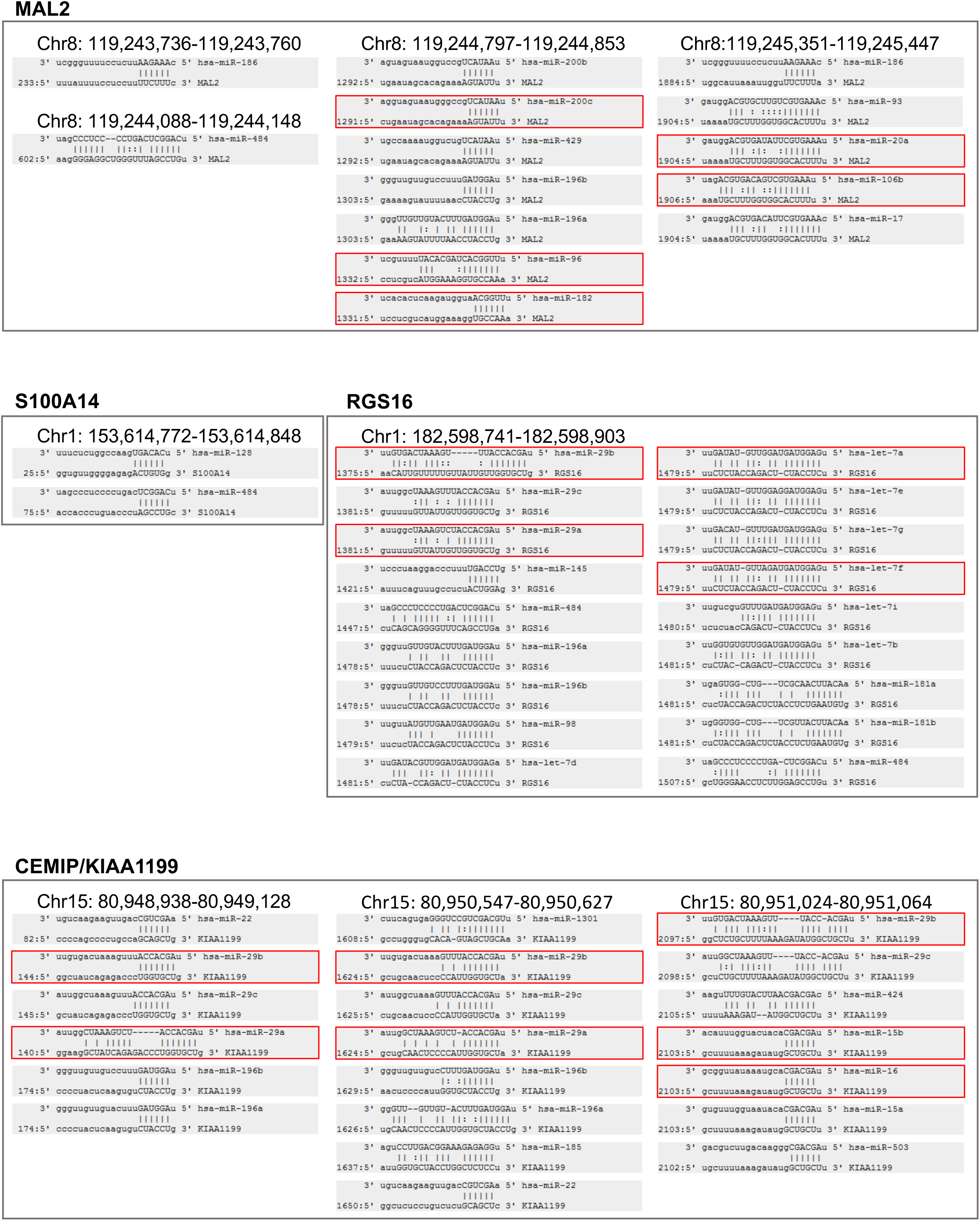
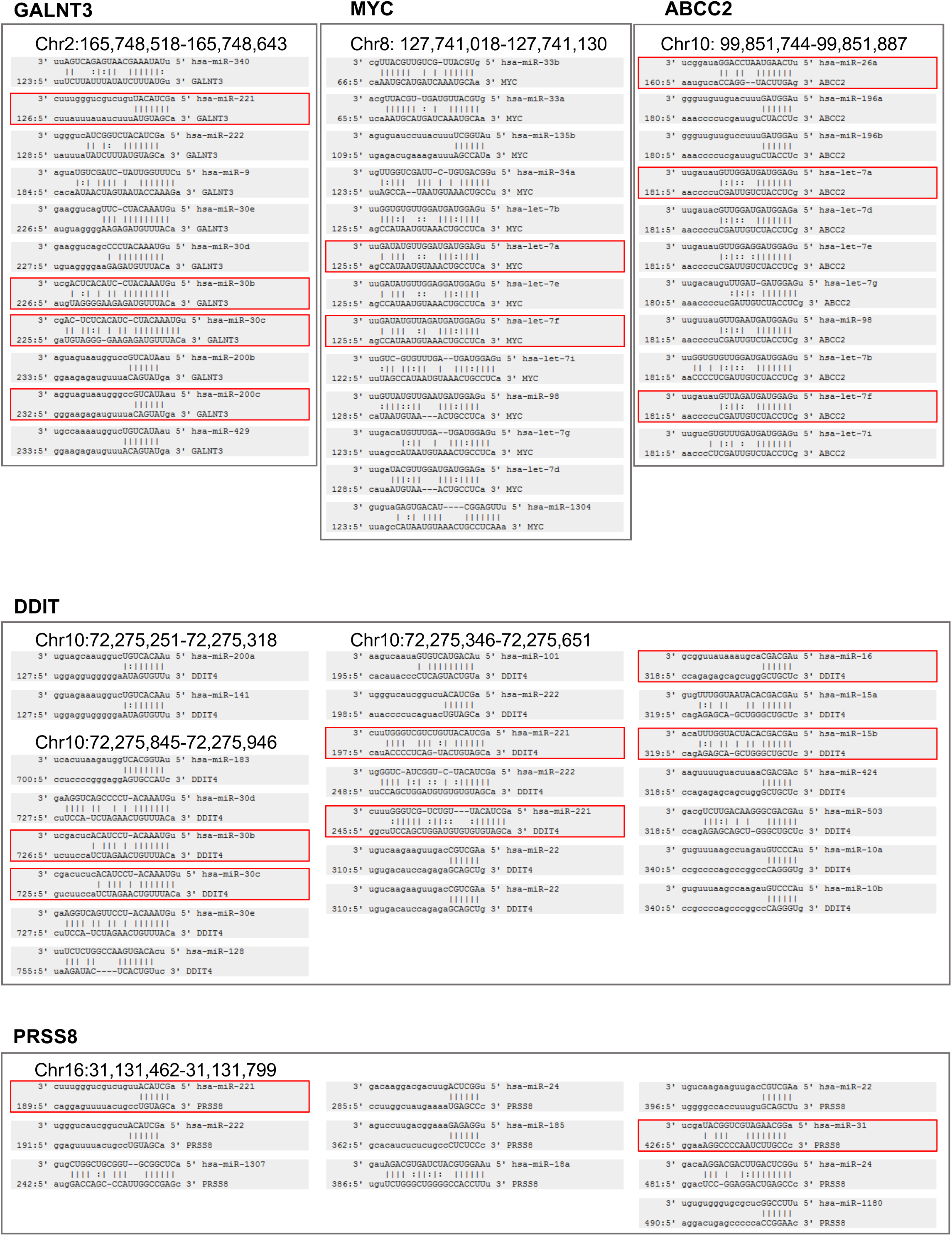
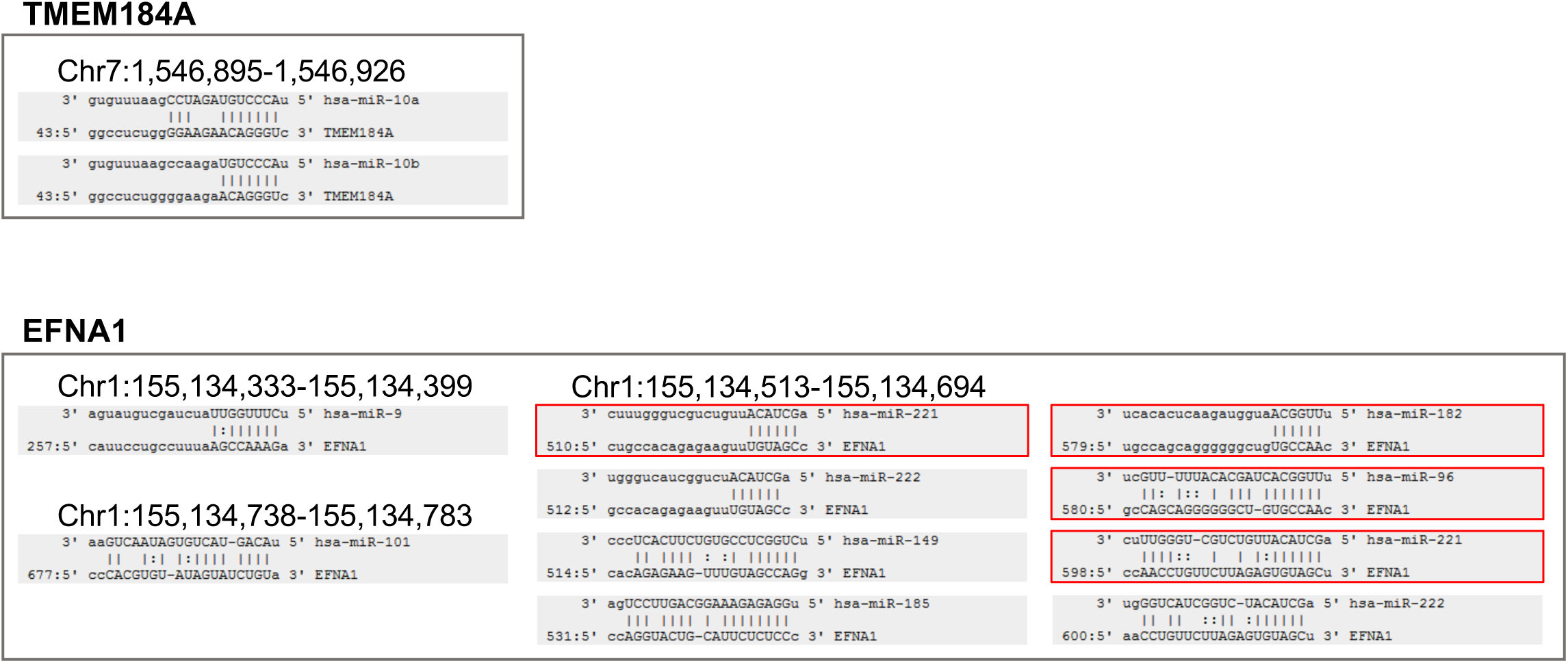
Prediction of miRNAs targeting to 3’-UTR regions of studied genes. Red outline – labeled top 25 expressed miRNAs in HCT116 cells.

**Supplemental Figure S7E (Related to Figure 7).**
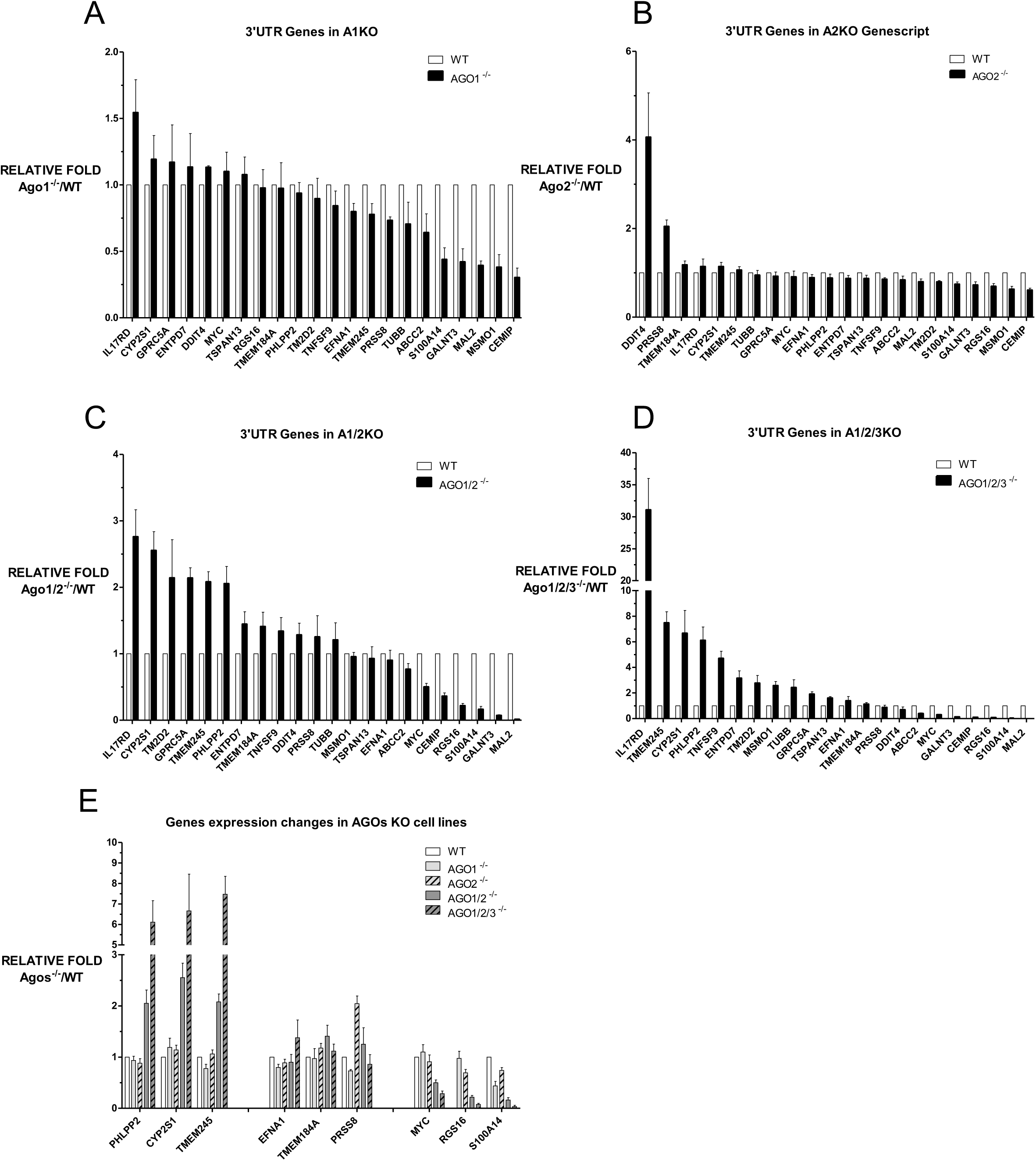
Expression level change of genes with AGO2 binding clusters located at 3’UTR measured by QPCR in AGO1^-/-^ cell line (A), AGO2^-/-^ cell line (B), AGO1/2^-/-^ cell line (C) and AGO1/2/3^-/-^ cell line (D). (E) Representative gene expression changes in AGOs knockout cel lines.

**Figure S7F (Related to Figure 7).**
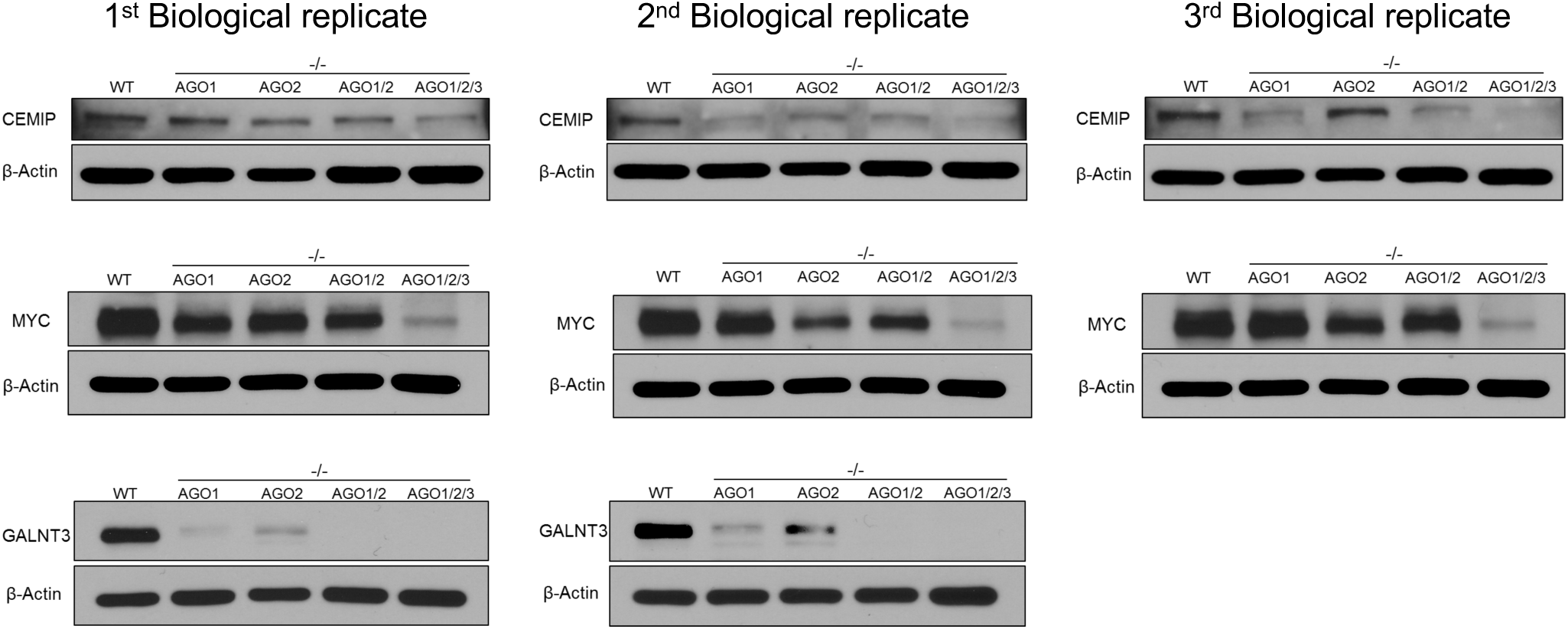
Biological replicates of Western-blots.

**Supplementary Figure S9.**
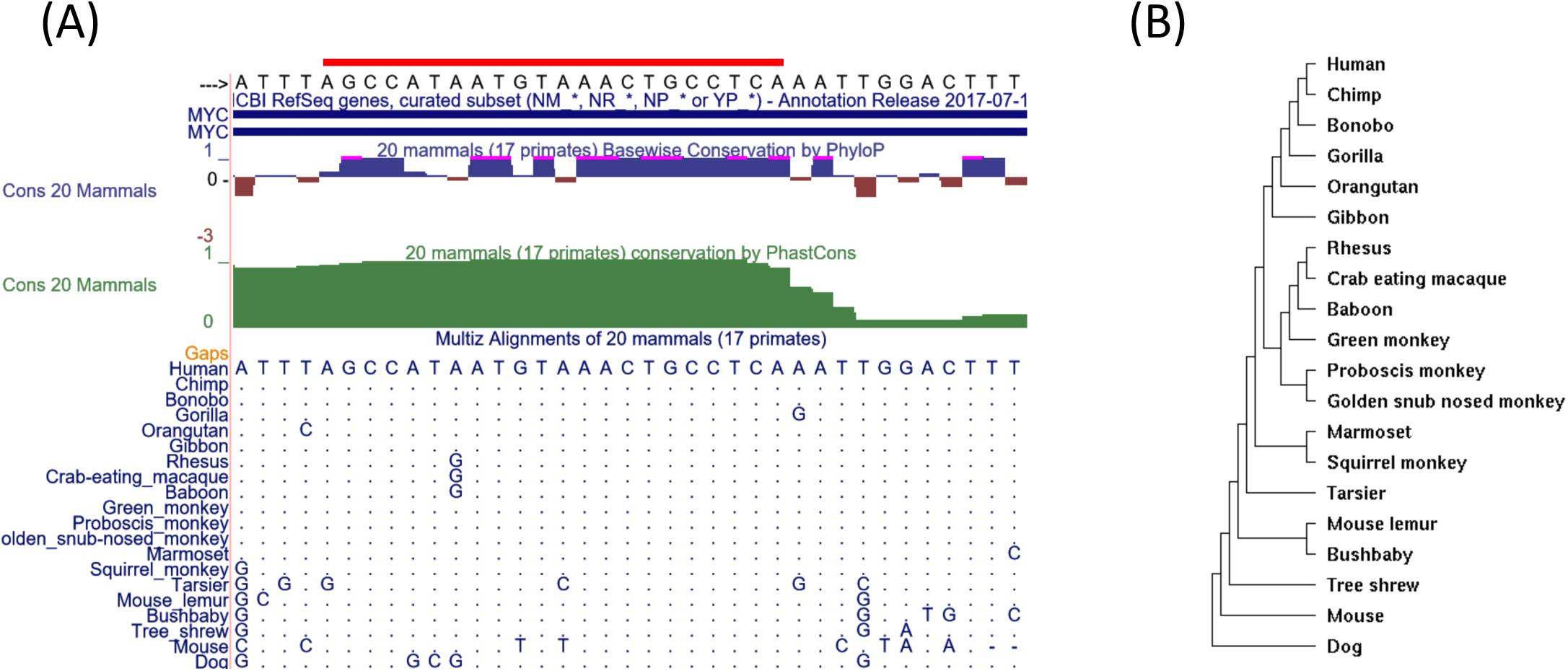
The sequence conservation analysis of a let-7 binding site within MYC 3’-UTR. (A) The binding site sequence alignments for 20 mammals. The PhyloP and PhastCons scores are also shown in the top part. (B) Phylogenetic tree model generated based on the sequence alignments for 20 mammals.

**Table S.**
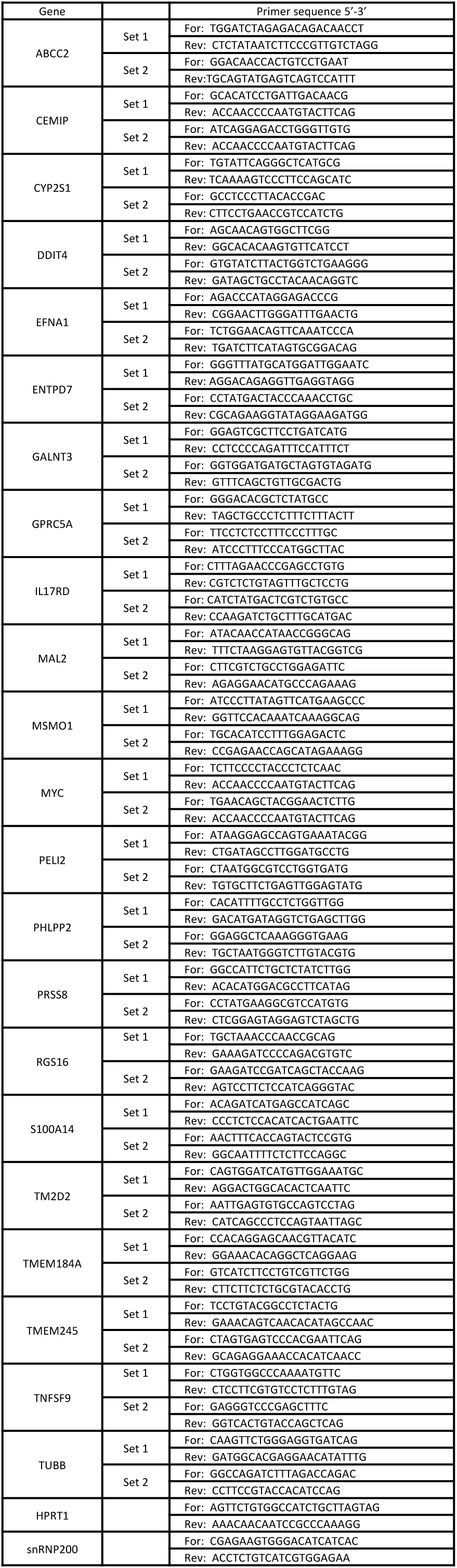
Primer sequences for validated genes.

## REFERENCES

1. Hammond, S.M. (2015). An overview of miRNAs. Advanced Drug Deliv. Rev. 87,3–14.

2. Bartel, D.P. (2018). Metazoan microRNAs. Cell 173, 20–51.

3. Xiao, Y and MacRae, I.J. (2019). Towards a comprehensive view of microRNA biology. Mol. Cell 75, 666–667.

4. Setten, R.L., Rossi, J.J., and Han, S. (2019). The current state and future directions of RNAi-based therapeutics. Nat. Rev. Drug Discov. 18, 421–446.

5. Shen, X. and Corey, D.R. (2018). Chemistry, mechanism, and clinical status of antisense oligonucleotides and duplex RNAs. Nucl. Acids Res. 46, 1584–1600.

6. Michlewski, G. and Caceres, J.F. (2019). Post-transcriptional control of miRNA biogenesis. RNA 25, 1–16.

7. Meister, G. (2013). Argonaute proteins: Functional insights and emerging roles. Nat. Rev. Genetics. 14, 447–459.

8. Yao, B., Li, S., and Chan, E.K.L. (2013). Function of GW182 and GW bodies in siRNA and miRNA pathways. Adv. Ex. Med. Biol. 768, 147–163.

9. Lewis, B.P., Shih, I., Jones-Rhoades, M.W., Bartel, D.P., Burge, C.B. (2003). Prediction of mammalian microRNA targets. Cell 115, 787–798.

10. Jackson, A.L., Burchard, J., Schelter, J., Chau, B.N., Cleary, M., Lim, L. and Linsley, P.S. (2006). Widespread siRNA “off-target” transcript silencing mediated by seed region sequence complementarity. RNA 12, 1179–1187.

11. Liu, J., Carmell, M.A., Rivas, F.V., Marsden, C.G., Thomson, J.M., Song, J., Hammond, S.M., Joshua-Tor, L. and Hannon, G.J. (2004). Argonaute2 is the catalytic engine of mammalian RNAi. Science 305, 1437–1441.

12. Meister, G., Lanthaler, M., Patkaniowska, A., Dorsett, Y., Teng, G. and Tuschl, T. (2004). Human argonaute2 mediates RNA cleavage targeted by miRNAs and siRNAs. Mol. Cell. 15, 185–197.

13. Su, S., Trombly, M.I., Chen, J., and Wang, X. (2009). Essential and overlapping functions for mammalian argonautes in microRNA silencing. Genes Devel. 23, 304–317.

14. Fire, A., Xu, S., Montgomery, M.K., Kostas, S.A., Driver, S.E., and Mello, C.C. (1998). Potent and specific genetic interference by double-stranded RNA in Caenorhabditis elegans. Nature 391, 806–811.

15. Zamore, P.D., Tuschl, T., Sharp, P.A. and Bartel, D.P. (2000). RNAi: double-stranded RNA directs the ATP-dependent cleavage of mRNA at 21 to 23 nucleotide intervals. Cell 101, 25–33.

16. Elbashir, S.M., Lendeckel, W., and Tuschl, T. (2001a). RNA interference is mediated by 21- and 22-nucleotide RNAs. Genes Dev. 15, 188–200.

17. Elbashir, S.M., Harborth, J., Lendeckel, W., Yalcin, A., Weber, K. and Tuschl, T. (2001b). Duplexes of 21-nucleotide RNAs mediate RNA interference in human cells. Nature 411, 494–498.

18. Corey, D.R. and Gagnon, K.T. (2019) Guidelines for Experiments using Double-Stranded RNAs and Antisense Oligonucleotides. Nucl. Acid Therapeutics 29, 116–122.

19. Capes-Davis A., Reid, Y.A., Kline, M.C., Storts, D.R., Strauss, E., Dirks, W.G., Drexler, H.G., MacLeod, R.A., Sykes, G. et al. (2012). Match criteria for human cell line authentication: Where do we draw the line? Int. J. Cancer. 132, 2510–2519.

20. Golden, R.J., Chen, B., Li, T., Braun, J., Manjunath, H., Chen, X., Wu, J., Schmid, V., Chang, T.C. et al. (2017). An Argonaute phosphorylation cycle promotes microRNA-mediated silencing. Nature 542, 197–202.

21. Gagnon, K.T., Li, L., Janowski, B.A. and Corey, D.R. (2014) Analysis of nuclear RNA interference in human cells by subcellular fractionation and argonaute loading. Nat. Protocols 9, 2045–2060.

22. Gagnon, K.T., Li, L., Chu, Y. Janowski, B.A., and Corey, D.R. (2014). RNAi factors are present and active in human cell nuclei. Cell Reports 6, 211–221.

23. Van Nostrand, E.L., Pratt, G.A., Shiskin, A.A., Gelboin-Burhart, C., Fang, M.Y., Sundararaman, B., Blue, S.M., Nguen, T.B., Surka, C. (2016). Robust transcriptome-wide discovery of RNA-binding protein sites with enhanced CLIP (eCLIP). Nat. Meth. 13, 508–514.

24. Ghandi, M., Huang, F.W., Jane-Valbuena, J., Kryukov, G.V., Lo, C.C., McDonald, E.R., Barretina, J., Gelfand, E.T., Bielski, C.M. et al. (2019) Next-generation characterization of the cancer cell line encyclopedia. Nature 569, 503–508.

25. Kalantari, R., Hicks, J.A., Li, L., Gagnon, K.T., Sridhara, V., Lemoff, A., Mirzaei, H., and Corey, D.R. (2016b). Stable association of RNAi machinery is conserved between the cytoplasm and nucleus of human cells. RNA 22, 1–14.

26. Chi, S.W., Zang, J.B., Mele, A., and Darnell, R.B. (2009). Argonaute HITS-CLIP decodes microRNA-mRNA interaction maps. Nature 460, 479–486.46.

27. Hafner, M., Landthaler, M., Burger, L., Khorshid, M., Hausser, J., Berninger, P., Rothballer, R., Ascano Jr. M., Jungkamp, A-C, Munschauer, M., et al. (2010). Transcriptome-wide identification of RNA-binding protein and microRNA target sites by PAR-CLIP. Cell 141, 129–141.

28. Erhard, F., Dolken, L., Jaskiewicz, L. and Zimmer, R. (2013). PARma: identification of microRNA target sites in AGO-PAR-CLIP data. Genome Biol. 14, R79.

29. Moore, M.J., Zhang, C., Gantmann, E.C., Mele, A., Darnell, J.C. and Darnell, R.B. (2014). Mapping argonaute and conventional RNA-binding protein interactions with RNA at single-nucleotide resolution using HITS-CLIP and CIMS analysis. Nat. Protocols 9, 263–293.

30. Bosson, A.D., Zamudio, J.R., and Sharp, P.A. (2014). Endogenous miRNA and target concentrations determine susceptibility to potential ceRNA competition. Mol. Cell 56, 347–359.

31. Broderick, J.A., Salomon, W.E., Ryder, S.P., Aronin, N. and Zamore, P.D. (2011) Argonaute protein identity and paring geometry determine cooperativity in mammalian RNA silencing. RNA 17, 1858–1869.

32. Elkayam, E., Faehlnle, C.R., Morales, M., Sun, J., and Joshua-Tor, L. (2017). Multivalent recruitment of human argonaute by GW182. Mol. Cell. 67, 646–658.

33. Hicks, J.A., Li, L., Matsui, M., Chu, Y., Volkov, O., Johnson, K.C., and Corey D.R. (2017). Human GW182 paralogs are the central organizers for RNA-mediated control of transcription. Cell Reports 20, 1543–1552.

34. Lewis, B.P., Shih, I-h., Jones-Rhoades, M.W., Bartel, D.P., and Burge, C.B. (2003). Prediction of mammalian microRNA targets. Cell 115, 787–798.

35. Grimson, A., Farh, K.K-H., Johnston, W.K., Garrett-Engele, P., Lim, L.P. and Bartel, D.P. (2007). MicroRNA targeting specificity in mammals, determinants beyond seed pairing. Mol. Cell 27, 91–105.

36. Friedman, R.C., Farh, K K-H., Burge, C.B., and Bartel, D.P. (2009). Most mammalian mRNAs are conserved targets for microRNAs. Genome Res. 19, 92–105.

37. Ruda, V.M., Chandwani, R., Algica, S., Bogorad, R.L., Akinc, A., Charisse, K., Tarkhovsky, A., Novobrantseva, T.I., and Koteliansky, V. (2014). The roles of individual mammalian argonautes in RNA interference in vivo. PLoS One 9, e101749.

38. Hafner, M., Landthaler, M., Burger, L., Khorshid, M., Hausser, J., Berninger, P., Rothballer, A., Asano, M., Jungkamp, A-C., Munschauer, M., Ulrich, A., Wardle, G.S., Dewell, S., Zavolan, M., and Tuschl, T. (2010). Transcriptome-wide identification of RNA binding protein and microRNA target sites by PAR-CLIP. Cell 141, 129–141.

39. Gosline, J.C., Gurtan, A.M., JnBaptise, C.K., Bosson, A., Milani, P., Dalin, S., Matthews, B.J., Yap, Y.S., Sharp, P.A., and Frainkel, E. (2016). Elucidating micrRNA regulatory networks using transcriptional, post-transcriptional, and histone modification measurements. Cell Rep. 14, 310–319.

40. Dang, C.V. (2012). MYC on the path to Cancer. Cell 149, 2012.

41. Kim, H.H., Kuwano, Y., Srikantan, S., Lee, E.K., Martindale, J.L., and Gorospe, M. (2009). HuR recruits let-7/RISC to repress c-Myc expression. Genes Dev. 23, 1743–1748.

42. Sampson, V.B., Rong, N.H., Han, J., Yang, Q., Aris, V., Soteropoulos, P., Petrelli, N.J., Dunn, S.P., and Krueger, L.J. (2007). MicroRNA Let-7a down-regulates MYC and reverts MYC-induced growth in Burkitt Lymphoma cells. Cancer Res. 67, 9762–9770.

43. Swier, L.J.Y.M., Dzikiewicz-Krawczyk, A., Winkle, M., van den Berg, A., and Kluiver, J. (2019). Intricate crosstalk between MYC and non-coding RNAs regulates hallmarks of cancer. Mol Oncol. 13, 26–45.

44. Cech, T.R. and Steitz, J.A. (2014). The noncoding RNA revolution – trashing old rules to forge new ones. Cell 157, 77–94

